# Programmable assembly of higher-order DNA nanostructures from microbial ssDNA staple libraries

**DOI:** 10.64898/2026.05.24.727530

**Authors:** Ming Hung Yen, Mallikarjuna Reddy Kesama, Yancheng Du, Jong Hyun Choi, Kevin V Solomon

**Affiliations:** Chemical & Biomolecular Engineering, University of Delaware, Newark, DE 19713; School of Mechanical Engineering, Purdue University, West Lafayette, IN 47907

**Keywords:** DNA origami, single-stranded DNA (ssDNA), DNA nanotechnology, Retron, Rolling circle replication, DNA staple libraries, Bio-enabled nanofabrication

## Abstract

Bottom-up manufacturing of structural DNA nanotechnology requires a long single-stranded DNA (ssDNA) scaffold and hundreds of short (∼30 nt) ssDNA staples. However, scaling production remains bottlenecked by the high economic cost and environmental footprint of solid-phase chemical staple synthesis. To address these limitations, a phage-free, biological nanomanufacturing platform engineered in *Escherichia coli* is developed here. Two intracellular strategies for producing programmable ssDNA were systematically evaluated: retron-based multicopy ssDNA (msDNA) synthesis via the Ec67 system and plasmid-encoded rolling circle replication (RCR). While native structural topology constraints within the retron (*msd*) cassette limit its sequence-design flexibility, the alternative RCR-driven engine successfully decouples ssDNA replication from sequence secondary structures, enabling the synthesis of arbitrary staples. This RCR platform reliably generates long circular ssDNA (cssDNA) precursors of at least 1.8 kb with exceptional sequence fidelity (>99%). Integrating programmable BseGI cleavage sites allows targeted strand-selective enzymatic processing to cleanly release stoichiometric, origami-grade pools of 32-nt staple strands. Atomic force microscopy (AFM) confirms that these biologically produced staples successfully drive the high-fidelity self-assembly of complex DNA tiles and hollow tubules. Crucially, robust structural folding is demonstrated directly within crude, unpurified cellular lysates, establishing a green, cost-effective framework for the one-pot fabrication of advanced DNA-based nanomaterials.

## INTRODUCTION

DNA origami exploits sequence complementarity to construct complex architectures with nanoscale precision[1-5]. The resulting DNA nanostructures have enabled a wide range of biological and biomedical applications, including biophysics, therapeutics, bio-sensing, and imaging[6-8]. In DNA origami, programmable and modular design is achieved by folding a long single-stranded DNA (ssDNA) *scaffold*, such as the 7,249-nucleotide (nt) M13mp18 genome, into desired geometries using a heterogeneous library of more than 150 unique, short ssDNA *staple* oligonucleotides. These staples provide design flexibility, structural predictability, and precise spatial addressability[9]. While DNA origami materials show excellent fault tolerance under atomic force microscopy (AFM), their functionalities rely on sequence accuracy and composition. As a result, translating DNA nanotechnology from laboratory-scale demonstrations to industrial applications requires scalable, cost-effective production of unique, defect-free ssDNA pools[10, 11].

Solid-phase chemical synthesis remains the gold standard for modern ssDNA manufacturing, but its stepwise phosphoramidite chemistry is highly resource-intensive, requiring a total 4,000 kg materials to produce 1 kg of ssDNA products[12]. Cumulative synthesis errors and purification losses further constrain achievable strand length and scale, inflating the cost of large, heterogeneous staple libraries[13, 14]. Enzymatic synthesis and amplification provide attractive aqueous workflows and access to kilobase-scale ssDNA; however, no universal, robust protocol has yet emerged[15]. In practice, yield, purity, and sequence compatibility depend strongly on template design, enzyme choice, reaction conditions, and downstream strand separation and purification workflows[16]. These constraints make reliable scale-up for diverse staple sets difficult[17]. /

In contrast, living systems naturally achieve low-cost, scalable ssDNA production during replication. Phage and phagemid platforms have been widely adapted for scaffold and, in some cases, staple strand generation; however, ssDNA generation in these systems is intrinsically coupled to coat protein expression and virion assembly[18-20]. As a result, ssDNA accumulates in viral particles and must be recovered by particle isolation and disassembly, complicating downstream processing[21]. Sequence design is further constrained by phage biology as engineered sequences must preserve essential genomic features required for coordinated protein expression and genome replication[22]. Realizing scalable ssDNA manufacturing therefore requires alternative, phage-free *in vivo* biomanufacturing strategies that decouple ssDNA synthesis from viral assembly while retaining full sequence programmability. In this context, *E. coli* provides an attractive platform due to its low-cost cultivation, genetic tractability, and compatibility with scalable bioreactor systems. However, existing *E. coli*-based approaches have generally produced modest, system-specific yields and have not resolved the combined challenges of sequence flexibility, production efficiency, and product recovery^11^.

Bacterial retron systems provide a well-established route for intracellular ssDNA production, and have been widely applied in *in vivo* gene editing[23, 24]. Retrons generate multicopy ssDNA-RNA hybrids (msDNA) via reverse transcription of a non-coding RNA template (Figure 1a). Engineering the *msd* region enables production of user-defined ssDNA[23]. We selected the Ec67 (Eco2) retron as a reference platform as it operates independently of RNase H and encodes an effector fused to the reverse transcriptase (RT), minimizing potentially deleterious interactions with host factors[25, 26]. Moreover, Ec67 is among the few retrons with reported absolute msDNA yields, enabling quantitative evaluation of production performance[27, 28]. Despite these advantages, engineering of Ec67 remains limited, and design rules for editable sequences within the *msd* are poorly defined[28]. An empirical assessment of Ec67 for programmable ssDNA production is therefore warranted.

**Figure 1.**
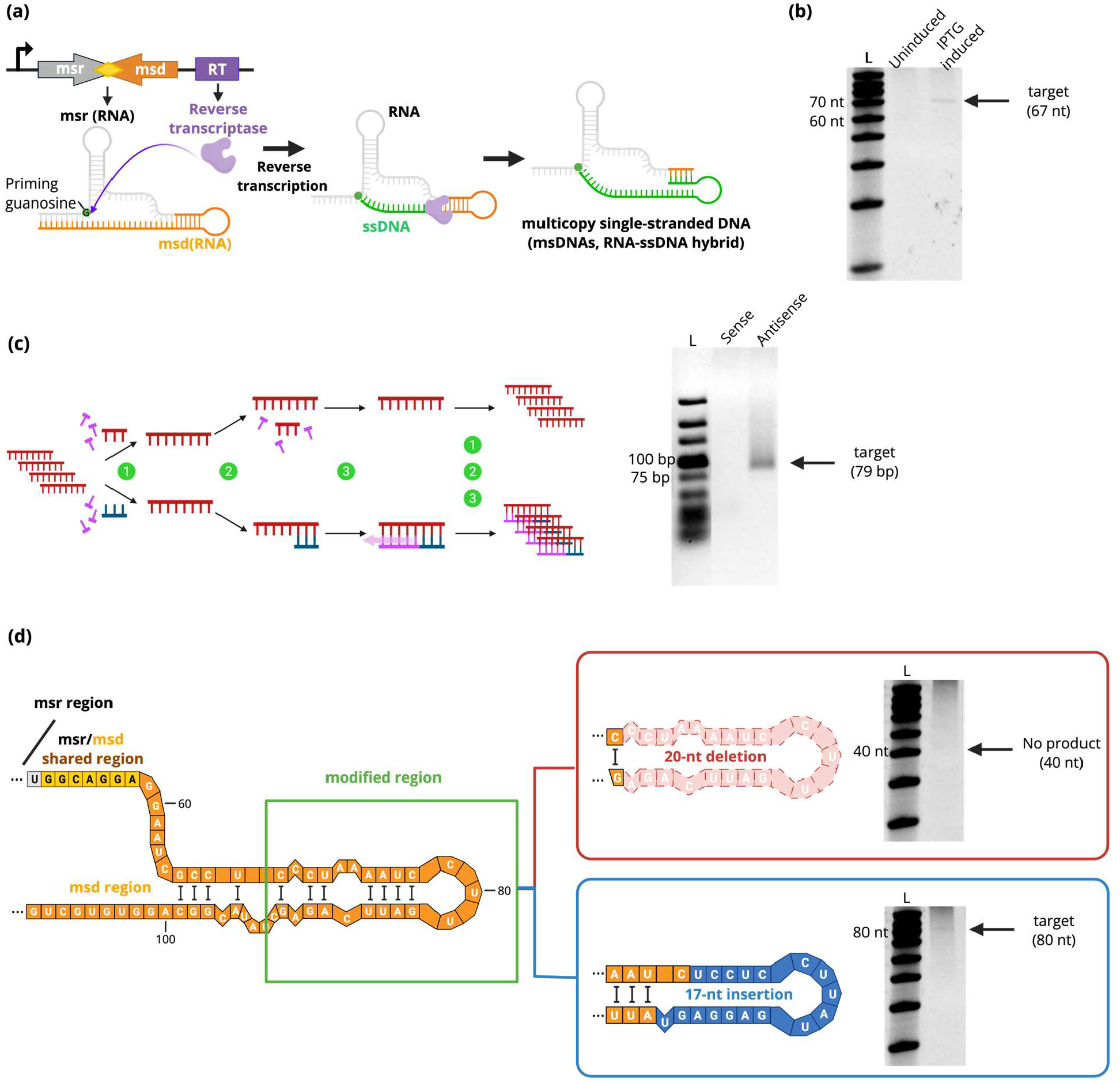
Characterization of retron Ec67 msDNA production, strand identity, and structural constraints. (a) Overview of retron architecture and msDNA biogenesis. (b) Ec67 msDNA production in *E. coli*. Denaturing PAGE of extracted DNA from uninduced and IPTG-induced cultures. (c) Strand identity confirmed by single-primer run-off reactions. Schematic (left) illustrates the single-primer run-off assay. The expected product (∼79 bp including primer overhang) forms exclusively when the antisense primer is used, as shown by the agarose gel (right), while no product is observed with the sense primer. (d) Structural constraints revealed by msd engineering. Empirically predicted secondary structure (left) of the WT Ec67 msr-msd RNA[62]. RNA structures of a 20-nt deletion (MHY-02) and 17-nt insertion (MHY-03) variants and msDNA production as shown by denaturing PAGE (right).

Plasmid-based rolling-circle-replication (RCR) has recently emerged as a complementary strategy for continuous *in vivo* ssDNA synthesis[29]. Unlike retrons, RCR uses a DNA template and relies on sequence-encoded initiation and termination elements, rather than RNA secondary structure, to control replication. Engineered RCR systems combining a dedicated replicase with defined origin sequences can generate circular ssDNA (cssDNA) corresponding to user-defined regions, with reported lengths up to ∼5 kb (Figure 2a)[24]. Co-expression of a single-stranded binding protein (SSB) stabilizes the product and improves yield. These long ssDNA molecules provide a versatile substrate for generating multiple staples through incorporation of sequence-encoded cleavage sites (e.g., restriction sites or catalytic DNA motifs)[30, 31]. However, prior RCR studies have primarily focused on continuous *in vivo* genome editing rather than on evaluation of ssDNA production capacity. Consequently, the practical performance of RCR-derived ssDNA as a programmable staple source remains an open question, motivating its systematic investigation in this work.

**Figure 2.**
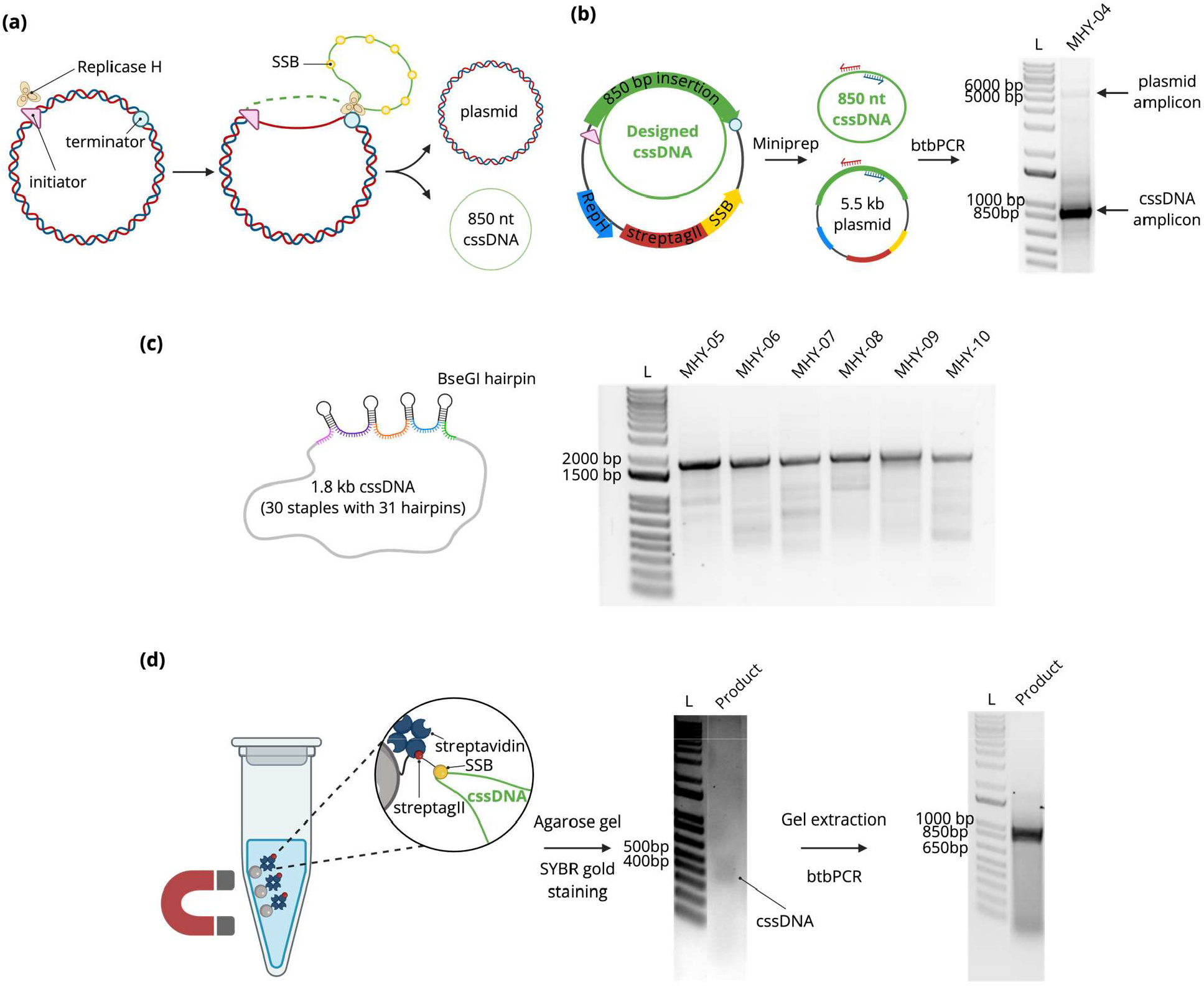
Evaluation of rolling-circle replication (RCR) as a platform for programmable long ssDNA synthesis. (a) Schematic overview of engineered RCR modules generating cssDNA corresponding to the cargo region. (b) Benchmarking cssDNA production with a 930-nt construct (MHY-04). Back-to-back PCR (btbPCR) using outward-facing primers selectively amplifies circular templates as shown by the agarose gel. (c) Visualization of robust expression of larger and complex cssDNA. Cargo length was doubled to ∼1.8 kb across six constructs, amplified by btbPCR, and shown by the agarose gel. (d) Direct visualization of cssDNA using bead extraction. StrepTagII-SSB fusion was captured by Streptavidin on the magnetic beads to enrich cssDNA. Sample was visualized by the agarose gel and SYBR Gold (middle), and btbPCR amplicon was visualized by the agarose gel (right).

In this work, we systematically evaluate retron-based ssDNA production using Ec67 and establish a complementary RCR platform for generating sequence-programmable ssDNA in *E. coli*. We demonstrate that retron output is strongly constrained by *msd*-dependent structural requirements, limiting its applicability for diverse sequence encoding. In contrast, the RCR system enables robust production of long cssDNA constructs (0.93–1.8 kb) with high sequence fidelity (>99%) and tolerance to repetitive and structure-forming sequences relevant to DNA origami design. By incorporating programmable cleavage motifs, these cssDNA precursors are enzymatically processed into balanced pools of short (32-nt) staple strands. We further demonstrate that bacterially produced staples support the assembly of higher-order DNA nanostructures, including tiles and nanotubules, with structural fidelity confirmed by AFM and performance comparable to chemically synthesized counterparts. Notably, DNA nanostructure assembly is maintained in crude cellular lysates, highlighting the potential for one-pot, minimally processed fabrication workflows. Taken together, this work establishes a phage-free, bio-enabled route for the scalable production of DNA origami staple libraries and demonstrates their direct applicability to nanostructure assembly. By decoupling ssDNA synthesis from viral assembly and enabling programmable, multi-strand encoding within single genetic constructs, this platform provides a foundation for sustainable, cost-effective, and potentially automated DNA nanomanufacturing.

## RESULTS AND DISCUSSION

### Retron Ec67 is sequence constrained for efficient ssDNA synthesis

We first evaluated retron-based ssDNA production using the Ec67 system as a model for intracellular synthesis. We cloned the Ec67 cassette into the pET28 expression vector (MHY-01) under a T7 promoter and expressed it in *E. coli*. Expression of the Ec67 cassette in *E. coli* produced detectable multicopy ssDNA (msDNA) only under induced conditions, with yields corresponding to ∼16 copies per cell (Figure 1b). To confirm strand identity and verify that the product was functional ssDNA, purified msDNA was subjected to single-primer run-off reactions (Figure 1c), which convert ssDNA templates into defined double-stranded products without exponential amplification. A product of the expected length (∼79 nt) was observed only when using the antisense primer, while the sense primer produced no detectable product. These results confirm the strand polarity and integrity of the Ec67-derived ssDNA (Figure 1c).

We next tested how flexible the retron platform is for encoding new sequences by introducing modifications within the *msd* region. While retrons vary widely in ssDNA performance, they share a conserved structure that may be used to inform engineering of Ec67[23]. Based on comparisons with engineered retrons such as Ec73, where portions of the *msd* stem–loop can be altered without loss of function, we hypothesized that a 20-nt segment in Ec67 might be dispensable[32-34]. To test this, we constructed a deletion variant (MHY-02). However, denaturing PAGE analysis detected no corresponding msDNA product, indicating that this region is required for efficient reverse transcription (Figure 1d).

To introduce sequence variation while preserving structural integrity, we designed a second variant (MHY-03) that maintains local base pairing. This construct included a 4-nt loop deletion and a 17-nt insertion that extends the stem while preserving predicted secondary structure. We modeled the RNA structure using the empirically-determined Ec67 architecture combined with minimum free energy predictions (Figure 1d, Figure S1)[34, 35]. Unlike the deletion variant, MHY-03 produced a detectable msDNA band upon induction, albeit at reduced yield (∼5 copies per cell), approximately one-third of the WT (Figure 1d, Table S4).

These results demonstrate that retron output is highly sensitive to RNA secondary structure, with efficient synthesis requiring preservation of native folding patterns. Mutations that disrupt the native structure markedly reduce or abolish yield, indicating the stringent structural dependence of the system. Consistent with this, comprehensive surveys across more than 100 retrons have reported similar structure-dependent limitations[36, 37]. As a result, retron-derived ssDNA has practically been restricted to relatively short (∼200 nt), structurally simple sequences[37, 38]. This strong sequence dependence limits the utility of retrons for generating diverse ssDNA libraries required for DNA origami, where large sets of arbitrary, sequence-specific strands are needed.

### Rolling circle replication enables sequence-programmable long circular ssDNA (cssDNA) production

To overcome the structural constraints inherent to retrons, we adapted RCR for intracellular ssDNA production. Unlike retrons, RCR uses a DNA template and is therefore largely decoupled from RNA folding constraints, offering greater sequence flexibility. We first benchmarked cssDNA production using a construct (MHY-04) encoding an ∼870 nt cargo containing 14 unique 32-nt staples flanked by 15 pairs of BseGI recognition sites, designed to form cleavable hairpins structures, and bounded by defined RCR initiator and terminator elements. Following IPTG induction, miniprepped samples were analyzed via back-to-back PCR (btbPCR) using outward-facing primers to selectively amplify circular templates. MHY-04-derived samples produced a strong amplicon at the expected size (930 nt) (Figure 2b). This signal persisted after linearization of the parent plasmid, while residual plasmid-derived amplification was eliminated, confirming that the product originated from cssDNA (Figure S2). We further optimized the btbPCR workflow by adding 10% (v/v) DMSO, which suppressed secondary structure formation from BseGI hairpins in the cssDNA template, increased target band intensity, and reduced nonspecific genomic amplification (Figure S3). We next increased cargo length and architectural complexity to ∼1.8 kb, distributing 30 unique staples and 62 identical BseGI recognition sites across six constructs (MHY-05 – MHY-10), collectively encoding the full 180-staple origami set. Despite the highly repetitive and structured sequence designs, all six constructs yielded robust, appropriately sized btbPCR amplicons, and next generation sequencing confirmed excellent fidelity (Figure 2c, Supplementary File). In all cases, polished consensus sequences matched the target design perfectly, suggesting that any minor read variances stem from technology-limited sequencing artifacts rather than synthesis errors[39].

To directly visualize and purify cssDNA, we used a single-stranded DNA-binding protein fused to a StrepTag II (SSB-StrepTagII) to capture ssDNA onto Strep-Tactin® XT magnetic beads followed by agarose gel analysis and SYBR Gold staining. Purified MHY-04 produced a single band at the expected size (930 nt ssDNA or ∼ 415 bp dsDNA) and yielded the correct btbPCR product and sequence (Figure 2d). For larger constructs (e.g. MHY-05, ∼1.8 kb), bead-purified material generated the expected btbPCR band and correct sequence but was not directly visible by SYBR Gold under our gel conditions (Figure S4). While this approach enabled enrichment of ssDNA, SSB binds DNA in a sequence-independent manner and likely dilutes recovery of the target cssDNA by co-enriching non-specific ssDNA species, resulting in the low yield observed in direct purification workflows. Attempts to directly sequence bead-purified material were limited by yield, suggesting that larger culture volumes will be required for direct recovery of preparative quantities of ssDNA. Nevertheless, these observations support the use of btbPCR as both a sensitive readout and a practical scale-up step for downstream processing.

### Enzymatic processing of cssDNA into defined staple libraries

We next evaluated whether microbially-derived cssDNA could be convered to staple libraries that drive higher-order DNA origami self-assembly. To generate sufficient single-stranded precursor, we performed scale-up btbPCR using a primer pair where one strand carried a 5’-phosphate modification. This modification selectively labels the undesired strand for degradation by Lambda exonuclease, enabling efficient recovery of the target ssDNA(Figure 3a). Following digestion, products were ethanol-precipitated to yield purified long ssDNA precursors.

**Figure 3.**
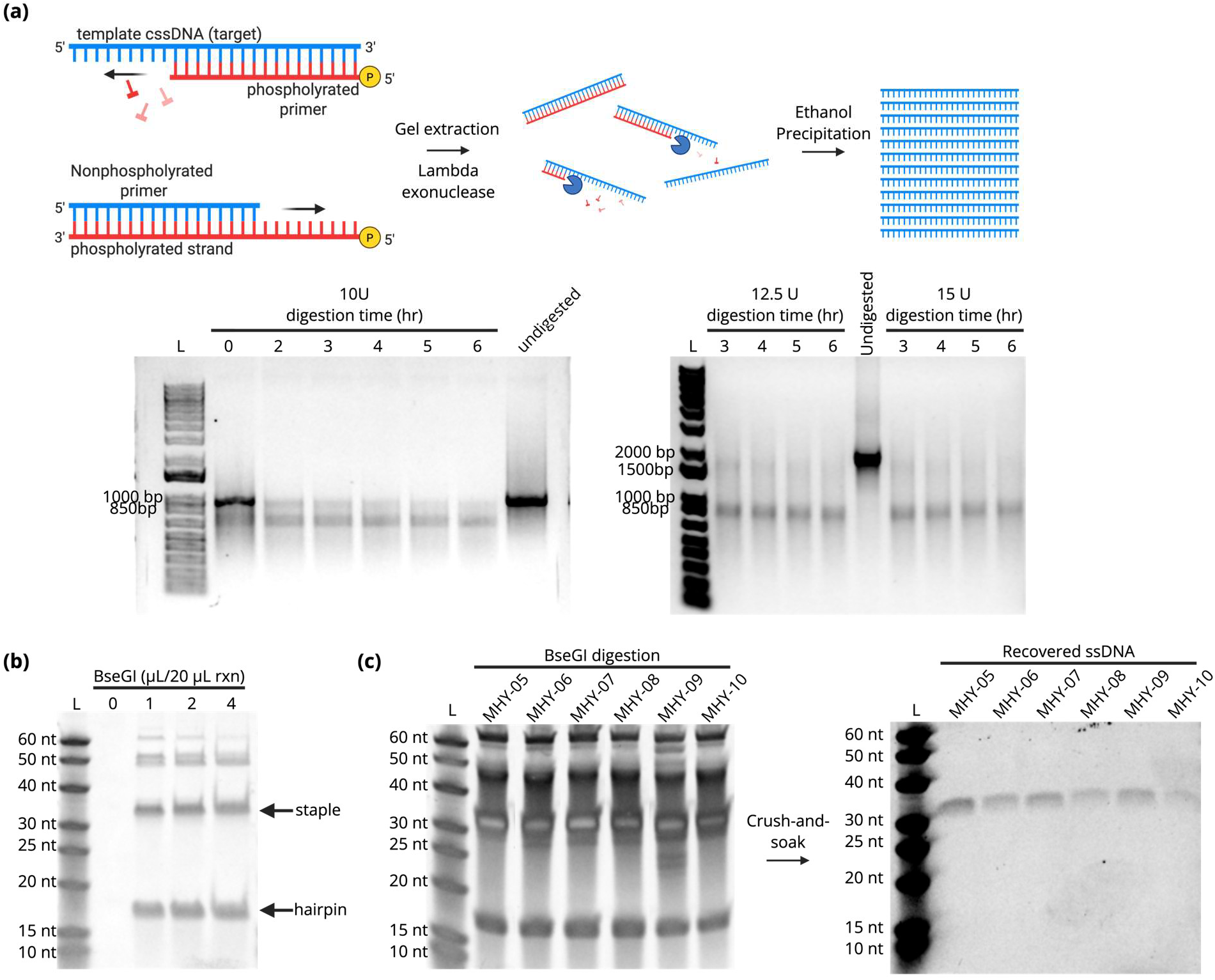
Preparation of ssDNA staples by strand-selective digestion and BseGI hairpin cleavage. (a) Schematic overview of btbPCR using a 5′-phosphorylated primer to selectively mark the unwanted strand for Lambda Exonuclease degradation. Time-course digestion assays identified near-complete strand removal for 930 bp (bottom left) and 1,800 bp (bottom right): 10 U enzyme for 930 bp amplicon (2 μg) at different time points (0, 2, 3, 4, 5, 6 hr) and undigested (lane 7). 12.5 U enzyme for 1,800 bp amplicon (1 μg) at different time points (3, 4, 5,6 hr), 15 U (1 μg) at different time points (3, 4, 5,6 hr), and undigested. (b) Time-course digestion assay identified BseGI cleavage of long ssDNA constructs: 1,800 nt cssDNA (200 ng) after 48 hr digestion with different enzyme concentrations (0, 1, 2, and 4 µL enzyme per 20 µL reaction). (c) Recovery and validation of pure 32-nt staples after BseGI cleavage using crush-and-soak strategy: 4 µL enzyme after 48 hr digestion with MHY-05-10 cssDNA samples. Crush-and-soak recovery for MHY-05-10 digested samples.

The purified ssDNA was heat-denatured and slowly cooled to promote intramolecular folding of the engineered BseGI hairpin structures required for cleavage. We then optimized BseGI enzyme loading and reaction time to maximize release of 32-nt staple strands (Figure 3b). To remove incomplete digestion products, the reactions were purified via denaturing PAGE and the desired products were recovered using a crush-and-soak workflow. In pilot experiments with model 32-nt ssDNA substrates, this method recovered ∼22% of target ssDNA (Figure S5). RCR-derived samples were processed identically and produced clean 32-nt staples across all constructs. Densitometry indicated staple yields in the hundreds of nanograms per construct, sufficient for downstream DNA origami assembly (Figure 3c). Together, these results establish a scalable enzymatic workflow for converting long, encoded ssDNA into defined staple libraries suitable for nanostructure fabrication.

### DNA nanostructure assembly using microbially produced staples

Next, we tested whether our microbially produced staples could successfully drive the self-assembly of complex DNA origami architectures. Using our optimized RCR platform in *E. coli*, we synthesized the180 unique core staples and 14 supplementary linker staples required to fold the target structures. We combined these 32-nt biological staples with a standard M13mp18 scaffold strand in 1x TAEM buffer, using the core pool to assemble planar rectangular tiles and adding the linker strands to drive tubule cyclization (Figure 4a)[40, 41]. We annealed the mixtures over a 16 hr thermal ramp before imaging. AFM confirmed the high-fidelity formation of well-defined, planar rectangular tiles featuring four distinct, sharp corners (Figure 4b, Figure S6). The assembled structures measured approximately 100 nm x 70 nm with a uniform height profile of ∼2 nm, perfectly matching the theoretical diameter of a single-layer DNA duplex[40]. Morphologically, we observed no discernible structural defects or structural differences between the origami tiles assembled with our RCR-derived staples and those built using commercial, chemically synthesized control oligonucleotides (Figure S7).

**Figure 4.**
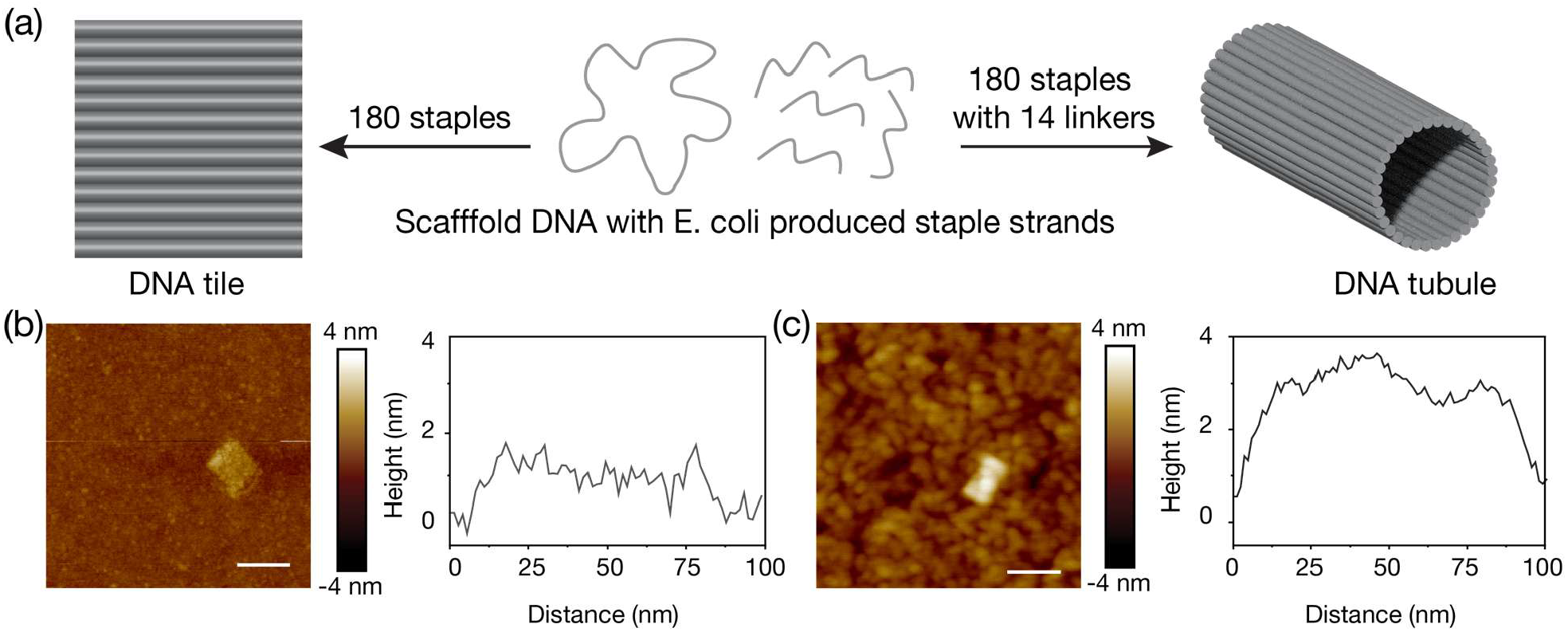
Construction of DNA tiles and tubules using the RCR-produced staples. (a) Schematic of DNA origami assembly. A long DNA scaffold binds with staple strands from E. coli, forming either a rectangular tile or hollow tubule. The unique 180 staples form a planar tile, while additional linkers induce a hollow tubule with a diameter of ∼30 nm in free solution. (b)-(c) AFM images and height profiles of DNA tile and tubule. The tile measures approximately 70 nm × 100 nm with a thickness of ∼2 nm. The tubule is half the size (70 nm × 50 nm) with twice the thickness (∼4 nm). Scale bars in AFM images: 100 nm.

The tiles can cyclize into tubules using linker staples that connect and seal the two opposite edges. The corners are not fully hybridized and are ready to bind with additional staples, which are seen as the single-stranded part in the AFM image (Figure 4b). With the linker strands from the RCR process, the one-pot thermal annealing generates hollow nanotubules. The AFM scan shows half the size of the tile with approximately 50 nm × 70 nm (Figure 4c). While the tubule is a 3D hollow structure with a diameter of approximately 30 nm, it becomes a 2D rectangle with a doubled thickness (∼4 nm) due to its maximal contact on the mica surface during AFM measurement. This observation confirms the accuracy of the strands from the RCR process in bacteria. While this study explores bacterial synthesis of oligo staples, it is also possible to program *E. coli* for production of custom scaffold sequences[42, 43].

Having established robust *in vivo* ssDNA manufacturing, we next asked whether we could assemble these complex nanostructures directly within crude biological matrices. That is, is there the potential for a one-pot synthesis of DNA origami with microbially derived staples. Previous studies reported that DNA origami can be assembled in the presence of living cells[44]. Here, we tested the feasibility of assembling DNA nanostructures across a gradient of raw *E. coli* cell lysates using our rectangular DNA tiles as a model system (Figure 5a). This challenge evaluated whether the target strands could successfully coordinate and fold amidst a crowded, heterogeneous background of genomic fragments, native proteins, cellular lipids, and the plasmids encoding the staple sequences themselves.

**Figure 5.**
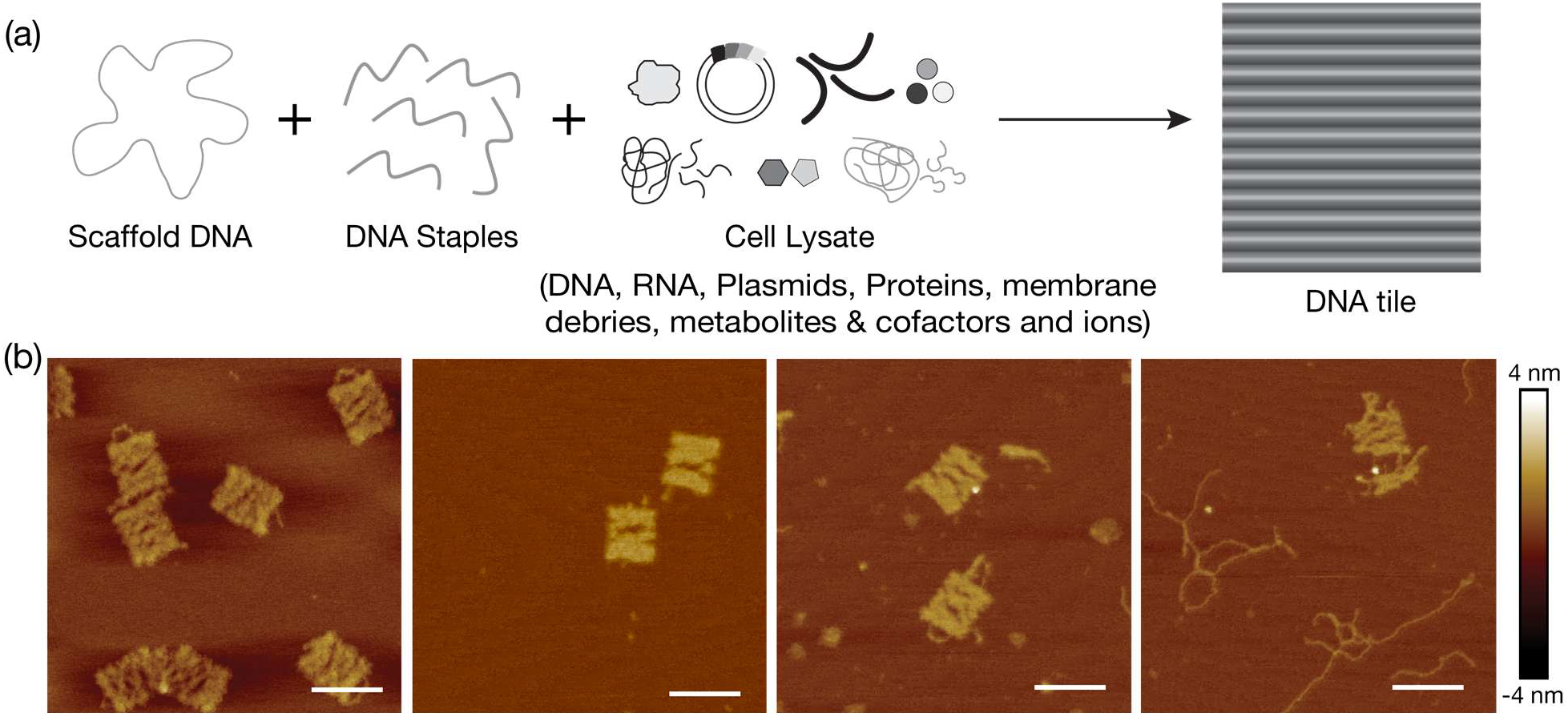
DNA origami assembly in a cell lysis environment. (a) Schematic illustrating DNA tile formation in the presence of cell components like plasmid encoding genes, genomic DNA (and fragments), RNA (and fragments), ribosomes, cellular proteins, membrane debris, lipids, metabolites, cofactors, and ATP, NAD+/NADH, and salt ions. (b) AFM images of DNA tiles assembled with 0, 1, 4, and 9% cell lysate (v/v). Scale bars in AFM images: 100 nm.

AFM imaging revealed successful formation of DNA rectangular tiles across a lysate concentration gradient of 0% - 9% (v/v) (Figure 5b). At lower lysate concentrations (≤ 1%), the AFM scans reveal sharp rectangles, proving that background cellular components do not inhibit precise scaffold routing or staple hybridization at these levels. At these concentrations, the cell lysate does not affect the scaffold folding and staple binding. At 4% lysate, small cellular fragments are observed along with 2D tiles. At 9%, there are clear signs of damage in DNA assemblies, with broken structures and long scaffold/plasmid strands observable. It is evident that cellular components like proteins, enzymes, and nucleic acids can disrupt DNA origami formation[45]. While nucleases digest DNA structures, plasmids can interfere with DNA self-assembly via competitive binding. Nevertheless, the persistence of intact tiles at moderate concentrations demonstrates that high-fidelity DNA nanotechnology fabrication remains entirely feasible within unpurified, biologically complex environments.

## CONCLUSION

In this work, we explored two distinct *in vivo* routes for sequence-defined ssDNA production in *E. coli*. The retron Ec67 system remained functional when the *msd* upper stem/loop was preserved but showed strong sequence-dependence and reduced yield upon modification. In contrast, plasmid-encoded RCR modules reliably produced at least 1.8 kb cssDNA. This template-driven engine successfully accommodated highly repetitive, staple-dense cargo that we cleanly processed into balanced 32-nt staple pools capable of driving higher-order DNA origami self-assembly. Together, these results provide a practical, directly accessible source of origami-grade staple ssDNA from standard *E. coli*.

Our findings align with and expand upon previous retron and RCR literature by mapping the precise topological design constraints of Ec67 and validating RCR for sequence-complex payloads. We show that Ec67 follows the same structural sequence design rules observed in other retrons (e.g. Ec73 and Ec86), where preservation of *msd* secondary structure dictates efficient msDNA production^17,24^. At the same time, we provide the first systematic evaluation of Ec67 tolerance to engineered sequence variation. For RCR, previous studies established kilobase-scale donor ssDNA (up to ∼5 kb) for genome editing, but did not examine tolerance to highly repetitive, hairpin-rich cargo^15^. Here, we demonstrate that RCR maintains exceptional sequence fidelity and functional output even in these demanding sequence contexts, extending its utility to origami-ready, structurally-complex payloads.

These capabilities suggest an accessible, sustainable origami manufacturing platform. Encoding large numbers of staples within a single long cssDNA enables tight control over staple stoichiometry, and preventing pool drift and imbalanced expression between cassettes. In principle, this approach could support integrated or one-pot assembly workflows, in which ssDNA production and nanostructure formation occur within a single system, building on prior demonstrations of intracellular DNA motif assembly[46]. Several technical limitations still remain. Our current cssDNA yields remain modest, and direct capture via downstream recovery is constrained by non-selective capture methods. The use of SSB, while effective for total ssDNA enrichment, lacks sequence specificity and likely co-isolates non-target ssDNA, reducing effective yield and purity. However, PCR amplification may be economically viable to circumvent these challenges under the right conditions. In addition, BseGI-mediated cleavage exhibits incomplete conversion, consistent with known sequence-context effects[47]. These limitations, however, appear tractable. RCR yields may be improved through engineering of initiation and termination elements, replicase expression, and growth conditions[48, 49], while sequence-specific capture strategies and programmable cleavage chemistries could improve recovery and processing efficiency. Future work may focus on a PCR-free pipeline that pairs higher yield with selective processing. For capture, sequence-specific strategies, such as transcription-regulatory proteins or short oligo hybridization with documented ssDNA selectivity, could enhance enrichment of target strands and improve purity over total SSB binding[50]. Similarly, DNAzyme-based cleavage systems could offer scarless, sequence-programmable cleavage and operate with biocompatible metal ions (e.g., Mg^2+^, Zn^2+^)[51].

Successful DNA origami formation validates our approach, highlighting sequence flexibility and accuracy. While this work focuses on staple oligonucleotides, scaffold strands with defined lengths may also be designed and produced. Beyond staple production, extension of this approach to scaffold synthesis and integration with isothermal or intracellular assembly strategies could further advance the development of fully biological DNA nanofabrication platforms. Future studies may thus be designed to suppress interference by cell components and promote self-folding and assembly under isothermal conditions[52-55]. Overall, this study integrates synthetic biology and DNA nanotechnology towards low-cost sustainable ssDNA production. Microbial engineering offers excellent programmability and sequence accuracy, laying a strong foundation for economic, large-scale manufacturing of designer DNA materials.

## ACKNOWLEDGMENTS

This material is based upon work supported by the National Science Foundation under Grant No. 2134603 to JHC and KVS.

## MATERIALS AND METHODS

### Bacterial Strains, Media, and Antibiotic Selection

*E. coli* DH5α was used for all plasmid cloning. One Shot™ BL21 Star™ (DE3) cells (ThermoScientific) were employed for all protein expression and ssDNA production, chosen for enhanced mRNA stability and low protease activity. Cultures were grown in Lysogeny Broth (LB, Miller, ThermoScientific). Antibiotics were added as required for plasmid maintenance: ampicillin (100 µg/mL), chloramphenicol (25 µg/mL), and kanamycin (50 µg/mL). Plasmids were transformed via heat-shock at 42 °C for 30 s, followed by 1 hr recovery in 200 µL SOC medium at 37 °C before plating on LB agar with antibiotic selection as needed.

### Cloning of plasmids

*E. coli* strains and plasmids used in this study are listed in Table S1. All molecular biology manipulations were carried out according to standard practices[56]. Plasmids were constructed via Gibson Assembly using NEBuilder® HiFi DNA Assembly Master Mix (New England Biolabs). Synthesized DNA inserts and primers used to generate precursors for assembly are detailed in Tables S2 -S3. DNA fragments used for assembly were amplified with Phusion High-Fidelity PCR Master Mix (Thermo), purified via gel and extracted using used Zymoclean Gel DNA Recovery Kit (Zymo). Plasmids were isolated and purified using E.Z.N.A.® Plasmid DNA Mini Kit (Omega). All kits were used according to the instructions of the manufacturer.

The retron Ec67 cassette was obtained from pLG006 (Addgene plasmid # 157884). MHY-01 was built using Gibson Assembly to clone the Ec67 cassette. Ec67 was digested from plG006 using XbaI and NdeI and assembled into the amplicon of pET28 (Primer 1 and 2). MHY-02 and MHY-03 were generated using site directed mutagenesis and the primers indicated in Table S2 (Primer 3-6)[57]. RNA secondary structures were predicted using empirical proposed structure and Vienna RNAfold Server (Vienna RNAfold Server (Version 2.6.3), May 19, 2026 used)[35, 58].

RCR constructs followed a pRC17-style design, with SSB-StrepTagII for cssDNA preservation and purification, RepH for cssDNA replication, and RCORI105-RCORI65 for replication module defining the RepH-amplified region[24]. gBlocks were assembled into pET28 and comprised (Table S3): (1) a P_LacIq_-driven SSB-StrepTagII cassette (P_LacIq_, iGEM RBS B0034, N-termial StrepTag II fused to *E. coli* SSB, PDB: 1EYG) (gBlock-1); (2) a T7-driven Replicase H (RepH) cassette (gBlock-2); and (3) an RCORI105-RCORI65 module with 14 staples and 30 BseGI sites cargo sequence (gBlock-3-5). Cargo sequences were cloned within the RCORI105-RCORI65 module for defined replication. CssDNA cargo design followed established strategies for enzymatic long ssDNA production, with each DNA origami DNA origami staple flanked by BseGI recognition sites[30]. The recognition site will adopt hairpin conformations after conversion to ssDNA and enable precise and programmable cleavage in ssDNA formats. The overall set of 194 staples (32-nt each) was based on the design of reported reconfiguration DNA origami[40]. An initial construct encoding 14 staples and 30 BseGI sites (∼930 bp cssDNA; MHY-04) was selected as a defined subset of the full origami design to bridge the tile into a tubular geometry for early validation of cssDNA production.

MHY-04 was constructed using Gibson Assembly with linearized pET28 plasmid (BglII and EcoRI digested) and gBlock-1-3. The remaining 180 staples were then distributed across six plasmids (∼1.8 kb cssDNA each; MHY-05-10) to maintain plasmid size below 10 kb for *E. coli* stability and cloning efficiency (MHY-05-10). All six constructs were built using Gibson Assembly with linearized MHY-04 PCR amplicon (Primer 9 and 10) and gBlocks (MHY-05: gBlock-6-8; MHY-06: gBlock-9-11; MHY-07: gBlock-12-14; MHY-08: gBlock-15-17; MHY-09: gBlock-18-20; MHY-10: gBlock-21-23).

### Sequencing and consensus generation

Plasmids and amplicons were all verified by Plasmidsaurus (whole-plasmid sequencing or amplicon sequencing). Both plasmids and btbPCR products for MHY-04–10 were sequenced by Plasmidsaurus on ONT R10.4.1 (Q20+) and assembled with their Super Accurate basecalling/polishing pipeline, yielding consensus accuracies often ≥Q60. Per-read accuracy and coverage metrics of btbPCR products for MHY-04–10 are reported in Supplementary File.

### Ethanol precipitation

Samples were mixing with prechilled 3X volumes of 100% ethanol, 1/10 volume of 3M Sodium Acetate (pH 5.2), and linear acrylamide (Invitrogen) to a final concentration 10–20 μg/mL. Product were incubated overnight at -20 °C, pelleted at 4 °C, 16,000*xg* for 30 min, washed twice with 70% ethanol.

### Electrophoresis & imaging

Agarose gels were pre-stained with SYBR safe (Invitrogen) and run in prechilled 1X TAE on ice; denaturing PAGE using 15% Mini-PROTEAN® TBE-Urea Precast Gels (Biorad) were run in pre-chilled 1X TBE and post-stained with SYBR gold (Invitrogen) for 10 min. All gels were imaged on Azure 400 Imager (Azure Biosystems). Densitometry was conducted in ImageJ.

### Retron ssDNA production and analysis

*E. coli* One Shot™ BL21 Star™ (DE3) carrying the respective retron constructs (MHY-01-03) were grown overnight at 37°C and 250 rpm, diluted 1:100, and cultured to mid-log phase (O.D._600_=0.4-0.6). Expression was induced with 0.1mM isopropyl β-d-1-thiogalactopyranoside (IPTG) and continued overnight expression at 37°C and 250rpm. Cells equivalent to 2 mL at O.D._600_ = 1 of overnight cultures were possessed by a single prep of miniprep and eluted in 50 µL nuclease-free water (Molecular Biology Grade, Fisher BioReagents). Miniprep eluates were then treated with RNase Cocktail™ Enzyme Mix (Invitrogen) for 30 min at 37°C and purified using the ssDNA/RNA Clean & Concentrator kit (Zymo). Purified msDNA products and 20/100 ssDNA ladder (IDT) were loaded with Novex™ TBE-Urea Sample Buffer (Invitrogen) and analyzed by denaturing PAGE. Gels were stained with SYBR Gold (Invitrogen) for 20 min, imaged on Azure 400 Imager (Azure Biosystems) with SYBR safe setting, and analyzed via ImageJ for band intensities. For strand validation, purified msDNA was subjected to single-primer PCR (run-off reaction) using either sense or antisense primer (Primer 7 and 8). Products were analyzed on 4% TBE-agarose gel alongside an Ultra Low Range DNA Ladder (Invitrogen), loaded with TrackIt™ Cyan/Yellow Loading Buffer and analyzed on imager. Given identical msr orientation and RT sequence across all variants, subsequent constructs were only analyzed by denaturing PAGE to assess ssDNA size and yield.

### RCR cssDNA Production and Analysis

*E. coli* One Shot™ BL21 Star™ (DE3) carrying RCR constructs (MHY-04-10) were grown overnight at 37°C and 250 rpm, diluted 1:100, and induced with 0.1mM IPTG at mid-log phase (O.D._600_ = 0.4-0.6). Overnight expression proceeded at 22°C and 250 rpm to favor RepH expression and cssDNA stability. Miniprep eluates from induced cell pellets were subjected to btbPCR using adjacent, outward-facing primers positioned centrally within the cargo region (primer 11 and 12). PCR reactions included 10% (v/v) DMSO, selected following optimization (Primer 13 and 14; Figure S3), with a 5.5 °C lower annealing temperature to traverse repetitive BseGI hairpins. Products were resolved by 1% agarose gels and verified by amplicon sequencing. For direct purification of cssDNA, induced cell pellets were lysed as previously described[59], and lysates were incubated with MagStrep® Strep-Tactin®XT beads (IBA) to capture SSB-Strep TagII bound cssDNA according to the manufacturer’s protocol. Bead-associated product was analyzed by agarose gels and SYBR Gold post-staining, gel-extracted, and further validated by btbPCR (Primer 15 and 16) and amplicon sequencing.

### Processing RCR-derived Long ssDNA into DNA Origami Staples

btbPCR was performed with one 5’-phosphorylated primer to selectively label the undesired strand (primer 11-P and primer 12, Table S2). Gel-purified btbPCR products were digested with Lambda Exonuclease (Thermo) at conditions determined by time-course agarose gels: 10 U for 6 h for ∼930 bp (2 µg DNA) and 15 U for 4 h for ∼1.8 kb (1 µg DNA). Reactions were heat-inactivated (80 °C, 15 min), ethanol-precipitated, and resuspended in 20 μL of DNase/RNase free water. Purified long ssDNA samples were heat-denatured. (95 °C, 5 min) and slow-cooled on the heating block (power off) to promote BseGI hairpin folding, then treated with FastDigest BseGI (Thermo) at conditions determined by time-course agarose gels: 4 µL (4 U) for 48 h (200 ng cssDNA) was adopted for subsequent preparations. 32-nt products were recovered by crush-and-soak with HiBind® DNA Mini Columns (Omega) as described[60, 61]. For benchmarking, commercial IDT 32-nt oligos were processed in parallel (Supplementary 5). For densitometry, 2% (v/v) of each purified sample was loaded on denaturing PAGE for analysis.

### Construction of DNA Origami Nanostructures

Rectangular tiles and hollow tubules were used in this study. The design of DNA origami, including scaffold folding pattern and staple arrangement, was reported elsewhere.[40, 41]. The sequence information is available in the supporting information (Table S5-S6). The M13mp18 scaffold was purchased from New England Biolabs. All other chemicals were purchased from Sigma-Aldrich. DNA nanostructures were synthesized by mixing the scaffold with staple DNA in 1x TAEM buffer (an aqueous solution of 40 mM trisaminomethane, 1 mM EDTA disodium salt, 20 mM acetic acid, and 10 mM MgCl_2_ at pH ∼8). The mixture was annealed in a Bio-Rad S1000 thermal cycler from 95 to 65 °C at 1 °C per 2 min, 65 to 60 °C at 1 °C per 25 min, 60 to 50 °C at 1 °C per 60 min, and 50 to 35 °C at 1 °C per 40 min, after which it was cooled down to 4 °C. The duration of the complete thermal cycle was 24 h.

### AFM Imaging of DNA Nanostructures

For AFM characterization, about 10 μL of DNA origami samples were pipetted onto a freshly cleaved mica surface and incubated for 5 min at room temperature in a closed Petri dish. After incubation, the mica was blown dry with compressed air, rinsed with ∼80 μL deionized (DI) water for about 3 sec, and then blown dry again with compressed air. AFM imaging was performed in air using the peak-force tapping mode with a Bruker Dimension Icon using Scanasyst-Air probes.

## TOC

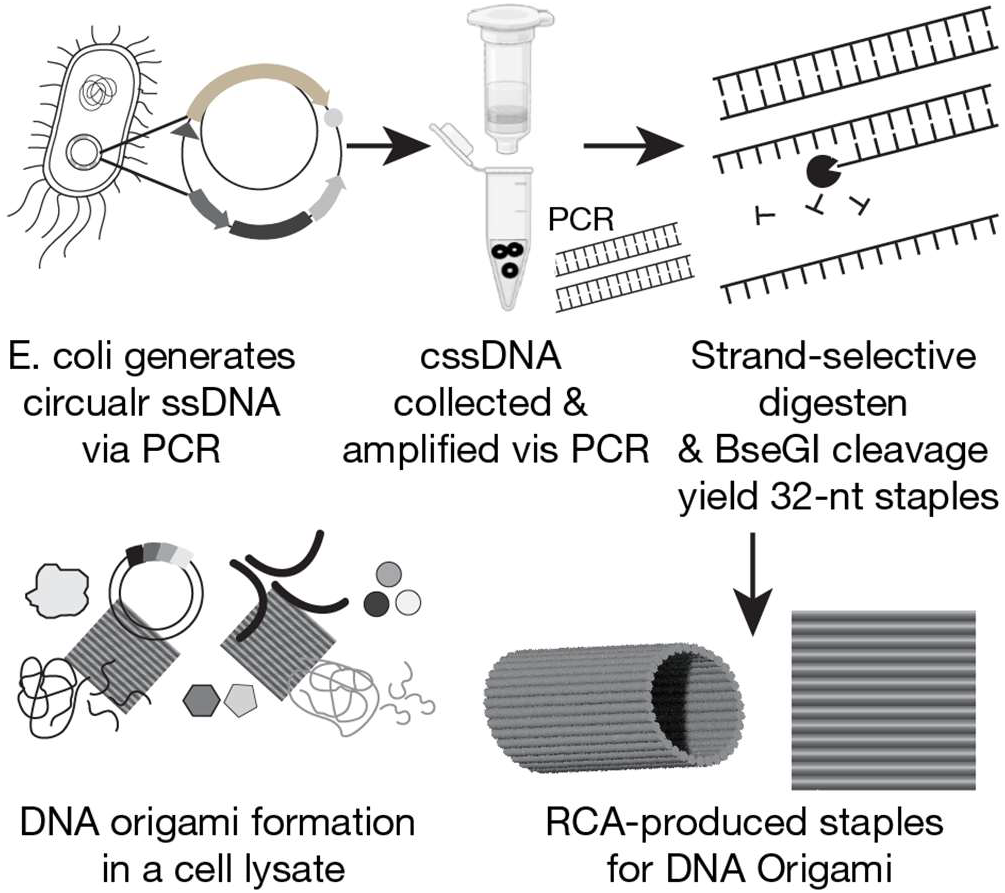

