## Supplementary Info for "Programmable assembly of higher-order DNA nanostructures from microbial ssDNA staple libraries"

**Table of Contents**

Figure S1. Ec67 retron RNA secondary structure.
Figure S2. Verification of btbPCR specificity for circular ssDNA templates
Figure S3. Optimization of DMSO concentration to improve btbPCR specificity and yield
Figure S4. Magnetic bead extraction and purification of MHY-05 cssDNA purification
Figure S5. Evaluation of crush-and-soak purification of 32-nt staples
Figure S6. DNA tubule formation using E. coli produced linker staples
Figure S7. DNA tile and tubule formation using IDT staples

Table S1. Strains and Plasmids
Table S2. Primers used in this study
Table S3. gBlock sequences
Table S4. Copy number of msDNA in Ec67-WT and Ec67-17-nt insertion
Table S5. Sequence information of 180 staples for the rectangular DNA origami tile
Table S6. Sequence information of 14 linker strands used for cyclization of a tile into a tubule

References

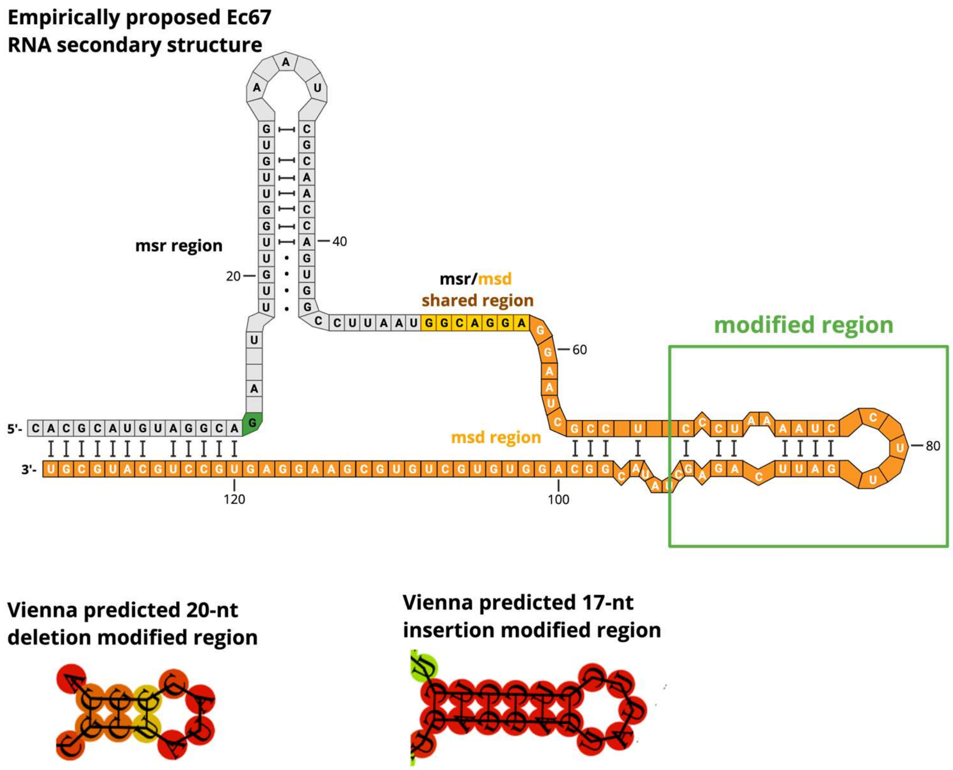


**Figure S1. Ec67 retron RNA secondary structure.** Empirically proposed Ec67 RNA secondary structure and Vienna predicted modified regions (Vienna RNAfold Server (Version 2.6.3), May 19, 2026)[1].

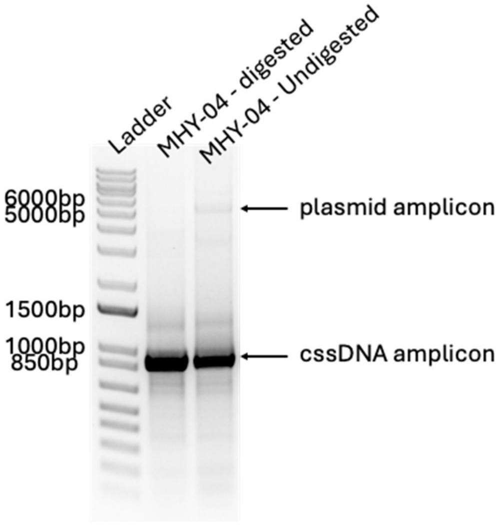


**Figure S2. Verification of btbPCR specificity for circular ssDNA templates.** Linearized samples lose plasmid-derived signal while retaining cssDNA amplification, confirming cssDNA-specific detection (primer 7 and 8 used).

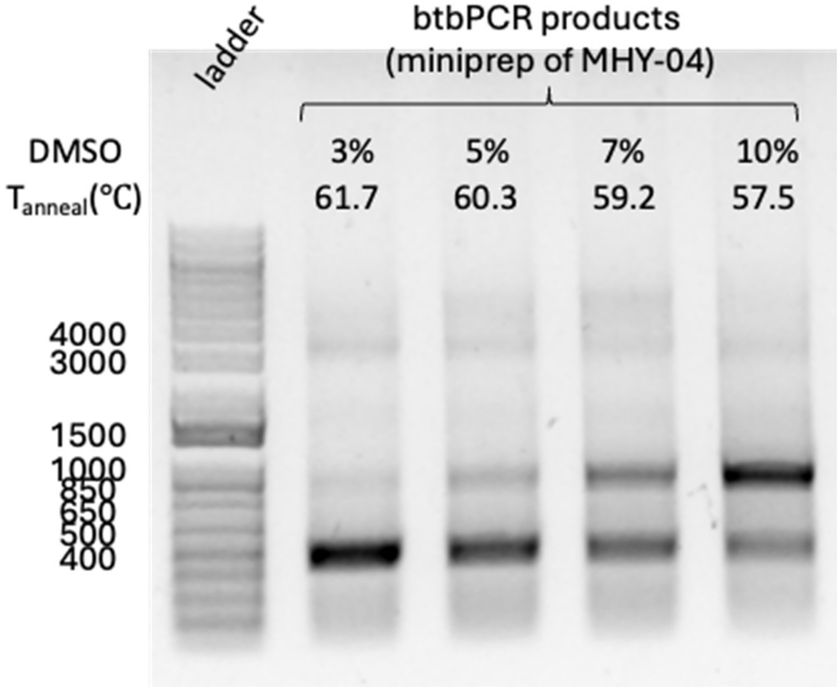


**Figure S3. Optimization of DMSO concentration to improve btbPCR specificity and yield.** Titration of DMSO (3-10%) during btbPCR showed that 10% yielded maximal amplification of the target cssDNA while minimizing nonspecific products (Primer 13 and 14). This condition was selected for all downstream experiments.

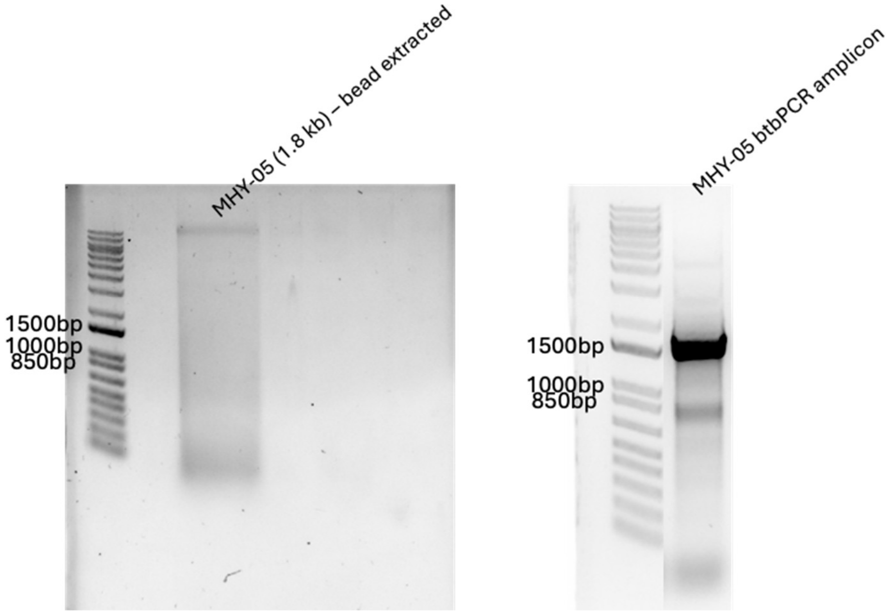


**Figure S4. Magnetic bead extraction and purification of MHY-05 cssDNA purification.** Bead-extracted cssDNA did not produce a strong or clearly visible band on agarose gel following SYBR Gold staining (left). However, btbPCR successfully amplified the bead-purified material, and the resulting amplicon was sequence-verified by amplicon sequencing, confirming correct cssDNA identity (Primer 15 and 16).

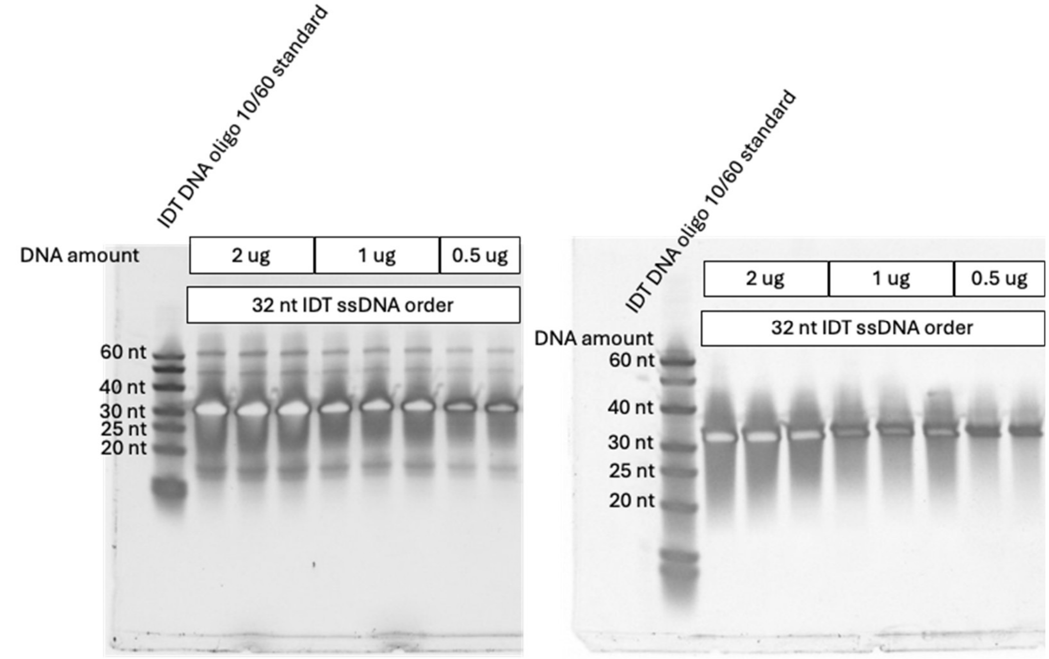


**Figure S5. Evaluation of crush-and-soak purification of 32-nt staples.** Denaturing PAGE analysis before and after crush-and-soak extraction shows recovery of a single, clean 32-nt band. The workflow was first benchmarked using synthetic 32-nt oligonucleotides, yielding ~22% recovery of pure ssDNA.

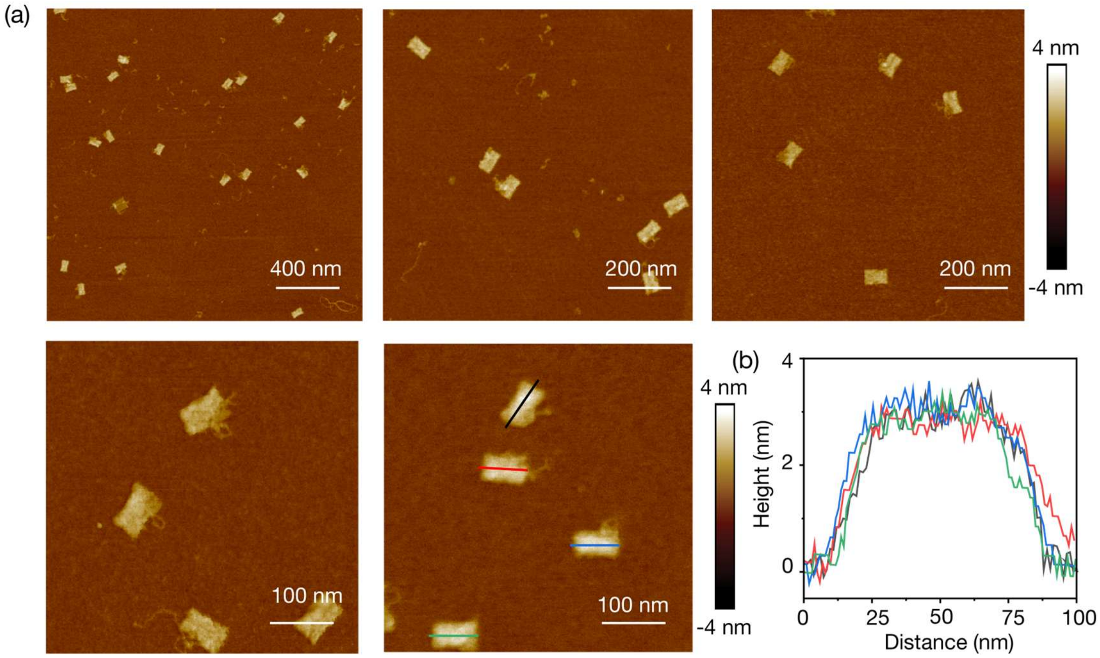


**Figure S6. DNA tubule formation using *E. coli*-produced linker staples.** (a) AFM images of DNA tubules formed via cyclization of DNA tiles using 14 linker strands from *E. coli*. Here, DNA tiles were first prepared with IDT staples and cyclized into tubules using 14 linker strands produced from *E. coli*. (b) Corresponding height profiles.

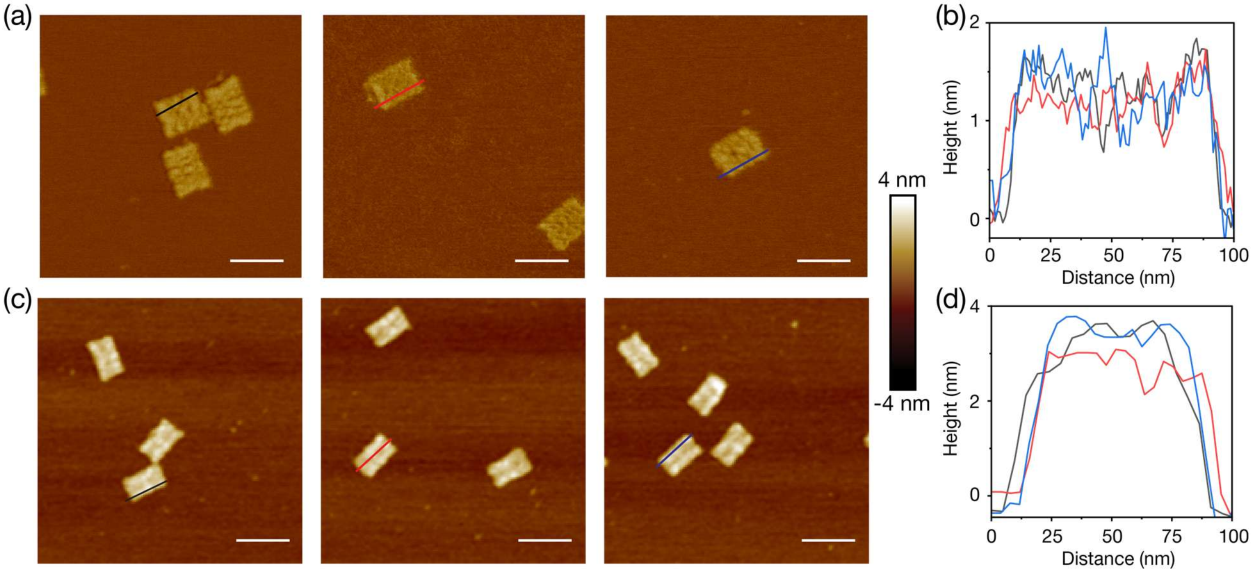


**Figure S7. DNA origami formation using *IDT* staples.** (a) AFM images and (b) height profiles of DNA tiles assembled using *IDT* staples. (c) AFM images and (d) height profiles of DNA tubules from *IDT* staples. Scale bars in AFM images: 100 nm

**Table S1.** Strains and Plasmids

| **Name** | **Genotype and relevant phenotype** | **Plasmid origin** | **Source** |
|---|---|---|---|
| **Strains** | | | |
| One Shot™ BL21 Star™ (DE3) | F–ompT hsdSB (rB–, mB–) gal dcm rne131 (DE3) | | |
| **Plasmids** | | | |
| pLG006 (Ec67 retron) | $Cm^R$, retron Ec67 cassette | P15A | Addgene (Plasmid #157884) |
| MHY-01 | $Kan^R$, IPTG inducible retron Ec67 | ColE1 | This study |
| MHY-02 | $Kan^R$, IPTG inducible retron Ec67 (20-nt deleteion) | ColE1 | This study |
| MHY-03 | $Kan^R$, IPTG inducible retron Ec67 (17-nt insert) | ColE1 | |
| MHY-04 | $Kan^R$, IPTG inducible RCR (staple-linker 1-14) | ColE1 | This study |
| MHY-05 | $Kan^R$, IPTG inducible RCR (staple 1-30) | ColE1 | This study |
| MHY-06 | $Kan^R$, IPTG inducible RCR (staple 31-60) | ColE1 | This study |
| MHY-07 | $Kan^R$, IPTG inducible RCR (staple 61-90) | ColE1 | This study |
| MHY-08 | $Kan^R$, IPTG inducible RCR (staple 91-120) | ColE1 | This study |
| MHY-09 | $Kan^R$, IPTG inducible RCR (staple 121-150) | ColE1 | This study |
| MHY-10 | $Kan^R$, IPTG inducible RCR (staple 151-180) | ColE1 | This study |

**Table S2.** Primers used in this study

| No. | Sequence 5' → 3' | Template | Final product |
|---|---|---|---|
| 1 | AGATATAAAGGAACATTATAAATTAATGTTA AATAGCTAAGTCGACAAGCTTGCGG | pET28 | MHY-01 |
| 2 | AACAAATCTGCCTACATGCGTGTTTATGAT GTATATCTCCTTCTTAAAGTTAAACAAAATT AT | pET28 | MHY-01 |
| 3 | CAGGAGGAATCGCCTCGCTATACG GCAGGTGTGCT | Ec67 cassette | MHY-02 |
| 4 | ACACCTGCCGTATAGCGAGGCGAT TCCTCCTGCC | Ec67 cassette | MHY-02 |
| 5 | TCGCCTCCCTAAAATCTCCTCCTTATG AGGAGTATTCAGAGCTATACGGCAGGT | Ec67 cassette | MHY-03 |
| 6 | CCGTATAGCTCTGAATACTCCTCATAAG GAGGAGATTTTAGGGAGGCGATTCCTC | Ec67 cassette | MHY-03 |
| 7 | CAATACAGATCTGGCAGGAGGAA TCGCCTC | Ec67 msDNA | Ec67 msDNA (79 bp, dsDNA) |
| 8 | AATAGAGTGTCTCCTTCGCACAGC ACACC | Ec67 msDNA | Ec67 msDNA (67 nt, ssDNA) |
| 9 | AATTCGAGCTCCGTCGACAAG | MHY-04 | MHY-05-10 |
| 10 | CCTGCAGGCATGCAAGCTT | MHY-04 | MHY-05-10 |
| 11 | AACGCACCAGTCATACCAATAACT TAAGGG | cssDNA (0.93-1.8 kb) | 930-1,800 bp amplicon |
| 12 | GCTACTGTGCAACCTGAAGGGGT | cssDNA (0.93-1.8 kb) | 930-1,800 bp amplicon |
| 11-P | /PHOS/AACGCACCAGTCATACCAAT AACTTAAGGG | cssDNA (0.93-1.8 kb) | 930-1,800 bp amplicon for Lambda Exo |
| 13 | GGTGGTTTTTCTTTTCACCAGTGAG ACTGCACAGTAGCCATTTCTAACG | cssDNA (930 nt) | 930 bp amplicon for DMSO testing |
| 14 | CGTTCCGCTATCGGCTGAATTTGA TTGACCTGAAGGGGTTGTTTTGG | cssDNA (930 nt) | 930 bp amplicon for DMSO testing |
| 15 | GCCTCGCCGGCAATAGTTACC CTTATTATCAAG | cssDNA (930 nt) | 930 bp amplicon for bead extracted btbPCR |
| 16 | CCTGAAGGGGTTGTTTTG GTGCATCCG | cssDNA (930 nt) | 930 bp amplicon for bead extracted btbPCR |

**Table S3.** gBlock sequences

| No. | Sequence 5' → 3' | Final product |
|---|---|---|
| 1 | CGTCCGGCGTAGAGGATCGATATAAACGCAGAAAGGCCCACCCGAA<br>GGTGAGCCAGTGTGACTCTAGTAGAGATCTACACTAGCACTATCAGA<br>GTTATTAAGCTACTAAAGCGTAGTTTTCGTCGTTTGCAGCCTGCCCG<br>CTTTCCAGTCGGGAAACCTGTCGTGCCAGCTGCAGGCGGGCTTTTCT<br>GTTCAGAACGGAATGTCATCATCAAAGTCCATCGGCGGCTCGTTAGA<br>CGGCGCTGCCGGAGCGGACTGCTGCGGGCGAGACTGCGCGCCGCC<br>GCTGAACTGATTGCCACCCTGCGGCTGCTGAGGCTGACCCCAACCG<br>CCCTGCGGCTGACCACCACCGATATTGCCACCTGCCGGAGCGCCAC<br>CACCCTGACGACCACCCAGCATCTGCATGGTGCCGCCAACGTTCAC<br>CACGACTTCTGTGGTGTAGCGATCCTGACCGGATTGATCGGTCCATT<br>TACGGGTACGCAGCTGACCTTCGATATAAACCTGAGAACCTTTACGC<br>AGATATTCGCTCGCCACTTCTGCCAGTTTGCCGAACAGCACAACGCG<br>GTGCCATTCAGTCTGTTCTTTCATCTCGCCGGTCGCTTTATCACGCCA<br>GGATTCGGAAGTAGCCAGCGTAATGTTGGCAACTGCGCCACCATTTG<br>GCATGTAGCGTACTTCCGGGTCCTGACCCAGATTACCAACGAGAATA<br>ACCTTGTTTACGCCTCTGCTGGCCATGGATCCTTTTTCGAACTGCGG<br>GTGGCTCCAGCTTGCCATATGGTACCTTTCTCCTCTTTCTCTAGTAAT<br>TCACCACCCTGAATTGACTCTCTTCCGGGCGCTATCATGCCATACCG<br>CGAAAGGTTTTGCACCATTCGATGGTGTCGAATATAGTCGACTCCTC<br>CTTTCG | MHY-04 |
| 2 | ATATAGTCGACTCCTCCTTTCGCTAGCAAAAAACCCCTCAAGACCCG<br>TTTAGAGGCCCCAAGGGGTTATGCTAGTTATTGCTCAGCGGTGGCAG<br>CAGTTTAGGTTAATTAAGCTGCGACTAGTAGACGAGTCCATGTGCTG<br>GCGTTCAAATTTCGCAGCAGCGGTTTCTTTACCAGACTCGAGTCAAA<br>GACTTTTTGATTTATTTATTAAATAATCACTATCTTTACCAGAATACTTA<br>GCCATTTCATATAATTCTTTATTATTATTTTGTCTTATTTTTTGAACTTG<br>AACTTGTGTTATTTCTGAAATGCCCGTTACATCACGCCATAAATCTAA<br>CCATTCTTGTTGGCTAATATAATATCTTTTATCTGTGAAATACGATTTA<br>TTTACTGCAATTAACACATGAAAATGAGGATTATAATCATCTCTTTTTT<br>TATTATATGTAATCTCTAACTTACGAACATATCCCTTTATAACACTACC<br>TACTTTTTTTCTCTTTATAAGTTTTCTAAAAGAATTATTATAACGTTTTA<br>TTTCATTTTCTAATTCATCACTCATTACATTAGGTGTAGTCAAAGTTAA<br>AAAGATAAACTCCTTTTTCTCTTGCTGCTTAATATATTGCATCATCAAA<br>GATAAACCCAATGCATCTTTTCTAGCTTTTCTCCAAGCACAGACAGGA<br>CAAAATCGATTTTTACAAGAATTAGCTTTATATAATTTCTGTTTTTCTAA<br>AGTTTTATCAGCTACAAAAGACAGAAATGTATTGCAATCTTCAACTAA<br>ATCCATTTGATTCTCTCCAATATGACGTTTAATAAATTTCTGAAATACT<br>TGATTTCTTTGTTTTTTCTCAGTATACTTTTCCATGTTATAACACATATG<br>TATATCTCCTTCTTAAAGTTAAACAAAATTATTTCTAGAGGGGAATTGT<br>TATCCGCTCACAATTCCCCTATAGTGAGTCGTATTAATTTCGCGGGAT<br>CGAGATCGATATTCTGTGGATAACCGTATTACCGCCTTTGAGTGAGC<br>TGATACCGCTCGCCGCAGCCGAACGACCGAGCGCAGCGAGTCAGTG<br>AGCGAGGAAGCGGAAGAGCGCCCAATACGCAAACCGCCTCTCCCCG<br>CGCGTTGGCCGATTCATTAATGCAGCTGGCACGACAGGTTTCCCGAC<br>TGGAAAGCGGGCAGTGAGCGCAACGCAATTAATGTGAGTTAGCTCA<br>CTCATTAGGCACCCCAGGCTTTACACTTTATGCTTCCGGCTCGTATGT | MHY-04 |

| | | |
|---|---|---|
| | TGTGTGGAATTGTGAGCGGATAACAATTTCACACAGGAAACAGCTAT<br>GACCATGATTACGCCAAGCTTGCATGCCTGCAGG | |
| 3 | CCAAGCTTGCATGCCTGCAGGTCGACTCTAGAGAAGGGTAACTAGC<br>CTCGCCGGCAATAGTTACCCTTATTATCAAGATAAGAAAGAAAAGGAT<br>TTTTGGATCCCCGGGTTTCATCCGCGCTTCGCGCGGATGAATAATAG<br>CCTCTTAATACTTTCATCTAAAATTGCATCCGCGCTTCGCGCGGATGC<br>AGGATTCTCGGCCTCCTGTTTAGTCTTGAGGTTCATCCGCGCTTCGC<br>GCGGATGAACTGTTTCTAATCCTAATCCTTCCTCCCGCTGACATCCG<br>CGCTTCGCGCGGATGTCGGTGGCAACGAGGACGCGCAGCACTGAG<br>CAAGCATCCGCGCTTCGCGCGGATGCTGACCCCCACTTATCCGCCT<br>GGTGTGACCGC | MHY-04 |
| 4 | CTGACCCCCACTTATCCGCCTGGTGTGACCGCCCCATCCGCGCTTC<br>GCGCGGATGGGTTAATTCCAGCGCCCTAGCGCCTCAACCCTTGCAT<br>CCGCGCTTCGCGCGGATGCACCTATTCCGGGCTATACTTATACGCTC<br>CTTGACATCCGCGCTTCGCGCGGATGTCGCTTTCTTCCCTTCCTTTC<br>TCGCGGTGATAAGCATCCGCGCTTCGCGCGGATGCTAACTATCTAAA<br>CCTCCTGAGTACCACGTTCCGCATCCGCGCTTCGCGCGGATGCGTT<br>CTTTTCCCCGTCAAGCTCTATCTGAGGG | MHY-04 |
| 5 | CGTTCTTTTCCCCGTCAAGCTCTATCTGAGGGCCCATCCGCGCTTCG<br>CGCGGATGGGCGTTGTTTCTGAGGGTGGCGGTAATCGGGGTCCATC<br>CGCGCTTCGCGCGGATGGACGGTTTTAGGGTTCCGATTTAGGCTCT<br>GAGACCATCCGCGCTTCGCGCGGATGGTGTTACGGAAAATGAGGGT<br>GGTGTGCTTTACCACATCCGCGCTTCGCGCGGATGTGATGGTTCGA<br>CCCCAAAAAACTTGTGGGCTTGTGCATCCGCGCTTCGCGCGGATGC<br>ACCAAAACAACCCCTTCAGGTTGCACAGTAGCCATTTCTAACGCACC<br>AGTCATACCAATAACTTAAGGGTAACTAGCCTCGCCGGCAATAGTTA<br>CCCTTATTATCAAGATAAGAAAGAAAAGGATTTTTCGCTACGCTCAAA<br>TCCTTTACAGAATTCGAGCTCCGTCGACAAGCTT | MHY-04 |
| 6 | AAGCTTGCATGCCTGCAGGTCGACTCTAGAGAAGGGTAACTAGCCTC<br>GCCGGCAATAGTTACCCTTATTATCAAGATAAGAAAGAAAAGGATTTT<br>TGGATCCCCGGGTAACATCCGCGCTTCGCGCGGATGTTCCTACGTC<br>GGGAGGTTCGCTAATAAATAATAACATCCGCGCTTCGCGCGGATGTT<br>CTCGATTCTTGATTTAAGGCTTAAATTAGGTTCATCCGCGCTTCGCGC<br>GGATGAATCTTTTTTTACTATTACCCCCTTATTTTGGAGCATCCGCGC<br>TTCGCGCGGATGCTATAATACCGCAGTATTGGACGCGGGTTATGATC<br>ATCCGCGCTTCGCGCGGATGATTTATGGTGAGGGGGATTCAATGGTA<br>ATTCCATCATCCGCGCTTCGCGCGGATGATGGCTGTTTTTATGTTATT<br>CTCTGCTATTGCAGCATCCGCGCTTCGCGCGGATGCTCTGGAAAGTT<br>AATTCTCCCGTCTATTGGGCAACATCCGCGCTTCGCGCGGATGTTTT<br>TATCCGTAGATTTTTCTTCCTAAAATCGATCATCCGCGCTTCGCGCGG<br>ATGATAGTGTTAATAATGTTGTTCCGTAGTTGAAAAACATCCGCGCTT<br>CGCGCGGATGTTTTGGCGCTAAATCTCAACTGATCTACTCGTTACAT<br>CCGCGCTTCGCGCGGATGTATTTCATTATTGCGTTTCCTCGGGATTG<br>TGA | MHY-05 |
| 7 | TATTTCATTATTGCGTTTCCTCGGGATTGTGAAACATCCGCGCTTCGC<br>GCGGATGTTAACTCAAATCATGCCAGTTCTTCTTTGCGTTGCATCCGC<br>GCTTCGCGCGGATGCATAAGGTAGGTCAGCCAGCCTATTTTCAAAGA<br>ACATCCGCGCTTCGCGCGGATGTTAAACCATATCCGGTTCTTGTCAT<br>TGACCGTGACATCCGCGCTTCGCGCGGATGTCTGGTGTAGCAGCTT<br>TGTTACGTCATGGAGCTGCATCCGCGCTTCGCGCGGATGCAAAATAA | MHY-05 |

| | | |
|---|---|---|
| | AATATTTACCTTCCCTACGGTGCTAACATCCGCGCTTCGCGCGGATG<br>TTCTTTTATGATAATTCACCTTTAGTTTCCGGAACATCCGCGCTTCGC<br>GCGGATGTTGGTCAGTGTATTCTTTTGCCTCTTACGTATAGCATCCGC<br>GCTTCGCGCGGATGCTGCGCCTTAATCGCTGGGGGTCATCATGAAA<br>CTCATCCGCGCTTCGCGCGGATGAGGTCGCGAATCTCCGTTGTACTT<br>GTAGCCGTGCCATCCGCGCTTCGCGCGGATGGCTATCGAGGGGGCT<br>ATGACCGAACTCTGGTG | |
| 8 | GCTATCGAGGGGGCTATGACCGAACTCTGGTGGGCATCCGCGCTTC<br>GCGCGGATGCCTTGCTATTTGATTATGAAAAGAGTGGTGGCAACATC<br>CGCGCTTCGCGCGGATGTTTACCCGCGTGGGCGATGGTTGTTGTTT<br>AAGTTCATCCGCGCTTCGCGCGGATGAAGTCTTTGCAAGCCTCAGCG<br>ACCTAAACCGACACATCCGCGCTTCGCGCGGATGTGCTACCCGACG<br>ATCCCGCAAAAGCCTTTTTTACCATCCGCGCTTCGCGCGGATGGTGG<br>TTCTTTTAATGAGGATTTATGGGGCATTGACATCCGCGCTTCGCGCG<br>GATGTCTGAGGGGTAAATTCAGAGACTGTCAAGGCATTCATCCGCGC<br>TTCGCGCGGATGAAATTCACTACAGAAAATTCATTTTCGTTACGTACA<br>TCCGCGCTTCGCGCGGATGTACAATTAGAAACTGTTGAAAGTTATGC<br>TACAAACATCCGCGCTTCGCGCGGATGTTGGAGATCTTTAGTTGTTC<br>CTTTGAAACTCATGCATCCGCGCTTCGCGCGGATGCACCAAAACAAC<br>CCCTTCAGGTTGCACAGTAGCCATTTCTAACGCACCAGTCATACCAA<br>TAACTTAAGGGTAACTAGCCTCGCCGGCAATAGTTACCCTTATTATCA<br>AGATAAGAAAGAAAAGGATTTTTCGCTACGCTCAAATCCTTTACAGAA<br>TTCGAGCTCCGTCGACAAG | MHY-05 |
| 9 | AAGCTTGCATGCCTGCAGGTCGACTCTAGAGAAGGGTAACTAGCCTC<br>GCCGGCAATAGTTACCCTTATTATCAAGATAAGAAAGAAAAGGATTTT<br>TGGATCCCCGGGTATCATCCGCGCTTCGCGCGGATGATAACCTTGTA<br>AAGTCTAATACTTTTGCCATCAACATCCGCGCTTCGCGCGGATGTTG<br>CCAACCAAAGGATTTAATACGTAATTCCGAGCATCCGCGCTTCGCGC<br>GGATGCTTGGCACTGCCTGGTTTCCGGCACGATCTTCGTCATCCGC<br>GCTTCGCGCGGATGACTGGGAACAGTTGCGCAGCCTGACCTCAAAC<br>CGCATCCGCGCTTCGCGCGGATGCGCCTTGCGCGAAGAGGCCCGC<br>ACCCCATCTAATCATCCGCGCTTCGCGCGGATGATCTGATACAAATA<br>ATTTTGATATCTCTGCGCAGCATCCGCGCTTCGCGCGGATGCTCCTT<br>CTTCTACGCAATTTCTTTCAATCAGGAGCATCCGCGCTTCGCGCGGA<br>TGCTGAGGCCAGGAAGGCCAGACGCGAAATGAGCCACATCCGCGCT<br>TCGCGCGGATGTGGCAGATGCTCACATTTAATGTTATTTTAACTGCAT<br>CCGCGCTTCGCGCGGATGCACCAACGGTTCCCACGGAGAATCATAT<br>TTGCTCCATCCGCGCTTCGCGCGGATGGATTTTGTGTTAAATGTAATT<br>AATCGTCTTAA | MHY-06 |
| 10 | GATTTTGTGTTAAATGTAATTAATCGTCTTAACGCATCCGCGCTTCGC<br>GCGGATGCGAATCCGTTTCCATTAAAAAAGGGGATTCTACACATCCG<br>CGCTTCGCGCGGATGTGATTTAACCAGACTCTCAGGCAATAGCTACC<br>TTCATCCGCGCTTCGCGCGGATGAAAATATTACGATTACCGTTCATCA<br>GAACGGTAACATCCGCGCTTCGCGCGGATGTTATACAAACCGGGGT<br>ACATATGAGACTGTCTGACATCCGCGCTTCGCGCGGATGTCTAAGCT<br>TAACCCAACCTAAGCCTAACGCCTCTCATCCGCGCTTCGCGCGGATG<br>AGGGAAAATGGATTTGCATCAGCAAAACCATTAGCATCCGCGCTTCG<br>CGCGGATGCTCTCCGGGCTTCTCCCGCAAAAGGATTTAGCCACATCC<br>GCGCTTCGCGCGGATGTGAATATCCTAAAAATTTTTATCCTAATTTTG | MHY-06 |

| | | |
|---|---|---|
| | GGCATCCGCGCTTCGCGCGGATGCCGGCCTTCTCAGGCATTGCATT TATTTATTG | |
| 11 | CCGGCCTTCTCAGGCATTGCATTTATTTATTGTACATCCGCGCTTCGC GCGGATGTATTTATCTACTGGTAAGAATTTGTCGTCTGGTTCATCCGC GCTTCGCGCGGATGAAATTTAGCAATTAAGCCCTACTGGTACTTTAAA CATCCGCGCTTCGCGCGGATGTTTATGCTTAATGGTCAAACTAAATAT ATGGAAGCATCCGCGCTTCGCGCGGATGCTAATTCTAAACAGGTTAT TGACCTAGTTGCACGCATCCGCGCTTCGCGCGGATGCGATGTTAATT TCAGCTCGCGCCCCCATTATATGTCATCCGCGCTTCGCGCGGATGAC AGAATTATTATTTTTCTTGTTCGGTAATGATACATCCGCGCTTCGCGC GGATGTATTCTCTGGTTTCTACATGCTCGCTTGCTTGATCATCCGCGC TTCGCGCGGATGATGAAACTTTCGCTTTGAAGCTCGTTCCTCTTTACA TCCGCGCTTCGCGCGGATGTATTTAAATCCTGACCTGTTGGAGGCTT CTGAGACATCCGCGCTTCGCGCGGATGTCAGCAATTATCAAAAGGAG CAATGATTTTTGTGCATCCGCGCTTCGCGCGGATGCACCAAAACAAC CCCTTCAGGTTGCACAGTAGCCATTTCTAACGCACCAGTCATACCAA TAACTTAAGGGTAACTAGCCTCGCCGGCAATAGTTACCCTTATTATCA AGATAAGAAAGAAAAGGATTTTTCGCTACGCTCAAATCCTTTACAGAA TTCGAGCTCCGTCGACAAG | MHY-06 |
| 12 | AAGCTTGCATGCCTGCAGGTCGACTCTAGAGAAGGGTAACTAGCCTC GCCGGCAATAGTTACCCTTATTATCAAGATAAGAAAGAAAAGGATTTT TGGATCCCCGGGTCACATCCGCGCTTCGCGCGGATGTGAAAACGTC TGTCGCTACTGATTCCCTCAATGCCATCCGCGCTTCGCGCGGATGGC TAATAATGGTTTCATTGGTGACATGAATAAGGCATCCGCGCTTCGCG CGGATGCCGGTGATATGGTAATGGTGCTACCTCAAGTCCGCATCCG CGCTTCGCGCGGATGCGGTTATGTTTAATGGAAACTTCCAAGATGAG TTCATCCGCGCTTCGCGCGGATGAACTCCCTAGTCCTCAAAGCCTCT TGTTTCGCTCCATCCGCGCTTCGCGCGGATGGAATATCATCAATGCT GGCGGCGGAATGCCGAATCATCCGCGCTTCGCGCGGATGATTCTGG CGGTGGCGGCTCTGAGGTGGCAAACAGCATCCGCGCTTCGCGCGG ATGCTGGAACGTGGCGGTTCTGAGGGTCTCTGGTTTTCATCCGCGCT TCGCGCGGATGAAATCCCACTCGAAAGCAAGCTGAGAATATATGTCA TCCGCGCTTCGCGCGGATGACTCCGCTAAGGCTCCTTTTGGAGCGG CCTTTCTCATCCGCGCTTCGCGCGGATGAGTCTCAGGTTCCGAAATA GGCAGTTGTTTGT | MHY-07 |
| 13 | AGTCTCAGGTTCCGAAATAGGCAGTTGTTTGTTTCATCCGCGCTTCG CGCGGATGAAACCCCGATACGGGCACTGTTACCGCTTTCCAGCATC CGCGCTTCGCGCGGATGCTCGACGGGTTAAAACTTATTACCGACGCT TACACATCCGCGCTTCGCGCGGATGTGGCGGTAGAGGGCTGTCTGT GGAGTTTAGCACCCATCCGCGCTTCGCGCGGATGGGTGGCGGAGTT TGTACTGGTGACCTATTCTCAGCATCCGCGCTTCGCGCGGATGCTGA TTCTCTCACTGATTATAAAAGCTGGCGGAGCATCCGCGCTTCGCGCG GATGCTCGTCAATGCGTGATGGACAGACCAAGGCCGGCCATCCGCG CTTCGCGCGGATGGCCGGCTTAATAGTGGACTCTTGTGGCTATTCGC CATCCGCGCTTCGCGCGGATGGCTCCCTTTTCGCCCTTTGACGTTGA TTTCGGCCCATCCGCGCTTCGCGCGGATGGGCACCTCACGTAGTGG GCCATCGCCTGCTGG | MHY-07 |
| 14 | GGCACCTCACGTAGTGGGCCATCGCCTGCTGGTACATCCGCGCTTC GCGCGGATGTAATATTGCTGCCAATGTAAATAATAGGGCTAATCATC CGCGCTTCGCGCGGATGATAGTTTGCCCTTTTATTACTGGTAGCCAT | MHY-07 |

| | | |
|---|---|---|
| | TCAACATCCGCGCTTCGCGCGGATGTTTTGATTCCCAATACGCAAAC CGCAGCTGGCTTCATCCGCGCTTCGCGCGGATGAACCACCATCTCA CTGGTGAAAAGCGGGCAGTCCCATCCGCGCTTCGCGCGGATGGGCA AACCCCAGGCGGTGAAGGGCTAGCTCACGACATCCGCGCTTCGCGC GGATGTCAGTTCGAGGCGGTGTTAATACTTATTTTAGTTCATCCGCG CTTCGCGCGGATGAAAAATATTTTGCTGCTGGCTCTCCTGTTGATGT CATCCGCGCTTCGCGCGGATGACGACAGGACGAATTCGAGCTCGGC ATGCAAGTCCATCCGCGCTTCGCGCGGATGGAGCGCAAATTTCACA CAGGAAACACGTCGTGGACATCCGCGCTTCGCGCGGATGTCATTAG GCGTATGTTGTGTGGAAAACTTAATTGCATCCGCGCTTCGCGCGGAT GCACCAAAACAACCCCTTCAGGTTGCACAGTAGCCATTTCTAACGCA CCAGTCATACCAATAACTTAAGGGTAACTAGCCTCGCCGGCAATAGT TACCCTTATTATCAAGATAAGAAAGAAAAGGATTTTTCGCTACGCTCA AATCCTTTACAGAATTCGAGCTCCGTCGACAAG | |
| 15 | AAGCTTGCATGCCTGCAGGTCGACTCTAGAGAAGGGTAACTAGCCTC GCCGGCAATAGTTACCCTTATTATCAAGATAAGAAAGAAAAGGATTTT TGGATCCCCGGGTATCATCCGCGCTTCGCGCGGATGATATGATAGG TGTTTATTCTTATTGGAGGTTACACATCCGCGCTTCGCGCGGATGTG GCTTTAACACGGTCGGTATTTCTTTACATAGCCATCCGCGCTTCGCG CGGATGGCGATTCTGTCAGAAGATGAAATTTTCTTTGTTTCATCCGCG CTTCGCGCGGATGAATGTATCCTGAGGCTTTATTGCTTTGCGTTGAT CATCCGCGCTTCGCGCGGATGATATAGCTTTGCCTTGCCTGTATGAA AATATAGACATCCGCGCTTCGCGCGGATGTCTATTGTATGTTGTTTAT TGTCGTATAACGCCGCATCCGCGCTTCGCGCGGATGCGATGGGATA CTTTACCTTTTGTCGTTGAGCGTTACATCCGCGCTTCGCGCGGATGT ATTGATTTATTACTGGCTCGAAATAAATATGCGCATCCGCGCTTCGCG CGGATGCGGTCTGGTCCAGACACCGTACTTATTTGCGAAGCATCCGC GCTTCGCGCGGATGCTCTCTAAACATGTTGAGCTACAGAAATGAAAG CCATCCGCGCTTCGCGCGGATGGCGTTCTTGCTTGCTATTGGGCGC AGGACTTA | MHY-08 |
| 16 | GCGTTCTTGCTTGCTATTGGGCGCAGGACTTAGGCATCCGCGCTTCG CGCGGATGCCCGCAAGATGAAAATAAAAACGGTAAATTAGGCCATCC GCGCTTCGCGCGGATGGCAACTAAGAGTGCGGTACTTGGTCAGCCG ATAGCATCCGCGCTTCGCGCGGATGCTAAACATTGATGCAATCCGCT TTTTTGCTTCCACATCCGCGCTTCGCGCGGATGTGACGATTGTCAGG GTAAAGACCTTAAAGGTAGCCATCCGCGCTTCGCGCGGATGGCTGC TATCTTATTTGGATTGGGAAACGCCTCAACATCCGCGCTTCGCGCGG ATGTTCCCTGTTTATTTTGTAACTGGCCAAAACCTCCCATCCGCGCTT CGCGCGGATGGGGTGTTCAGACGCTCGTTAGCGTGCAAAATAAACA TCCGCGCTTCGCGCGGATGTTTATTAAGTCGTCTGGTAAACGATATC CAGTAGCATCCGCGCTTCGCGCGGATGCTACCTGTGCTCTTACTATG CCTCAATATTTA | MHY-08 |
| 17 | CTACCTGTGCTCTTACTATGCCTCAATATTTACACATCCGCGCTTCGC GCGGATGTGGTAACTGGGCTTCGGTAAGATACTGTAAAGGACATCCG CGCTTCGCGCGGATGTCCGTTATTGTTTCTTGCTCTTATTAATGCGCC CCATCCGCGCTTCGCGCGGATGGGAGTCTTATTCTTGTGGGTTATCT TTGTTCAAGCATCCGCGCTTCGCGCGGATGCTTGATGAAATTCACAA TGATTAATAGTTCGTACCATCCGCGCTTCGCGCGGATGGTAATGAAT CTCAAGCCCAATTTAGAATCTTTCGCATCCGCGCTTCGCGCGGATGC GGTTGAAATATGAATTTTCTATTTTTCCTTCAACATCCGCGCTTCGCG | MHY-08 |

| | | |
|---|---|---|
| | CGGATGTTTCCGTCACTTATTCCGTGGTGTTTGGGTATCCCATCCGC<br>GCTTCGCGCGGATGGGTGACGGTATGTTGCCACCTTTACGTAATAAC<br>ACATCCGCGCTTCGCGCGGATGTGTTTTAGTTCGGTTCCCTTATGAA<br>GATTACTGCCATCCGCGCTTCGCGCGGATGGCTTGGTACGTTCCGG<br>CTAAGTAATGATTTGGTGCATCCGCGCTTCGCGCGGATGCACCAAAA<br>CAACCCCTTCAGGTTGCACAGTAGCCATTTCTAACGCACCAGTCATA<br>CCAATAACTTAAGGGTAACTAGCCTCGCCGGCAATAGTTACCCTTATT<br>ATCAAGATAAGAAAGAAAAGGATTTTTCGCTACGCTCAAATCCTTTAC<br>AGAATTCGAGCTCCGTCGACAAG | |
| 18 | AAGCTTGCATGCCTGCAGGTCGACTCTAGAGAAGGGTAACTAGCCTC<br>GCCGGCAATAGTTACCCTTATTATCAAGATAAGAAAGAAAAGGATTTT<br>TGGATCCCCGGGTCTCATCCGCGCTTCGCGCGGATGAGACGATTTTT<br>TCCTGTTGCAATGACACTTCTCACATCCGCGCTTCGCGCGGATGTGG<br>TGAATTTCTGGATATTACCAGTCTTTTACTGCATCCGCGCTTCGCGCG<br>GATGCAGAATGTAGTTCTTCTACTCAGGGCTACAACGGCATCCGCGC<br>TTCGCGCGGATGCCCTGGCGTATAAGGGATTTTGCCGGAGTCCAAC<br>CATCCGCGCTTCGCGCGGATGGTTGCCCGTCAAACAGGATTTTCGC<br>CCTGATATGCATCCGCGCTTCGCGCGGATGCACCTCTGTTTAATGGC<br>GATGTTTTCCATTTCGTCATCCGCGCTTCGCGCGGATGACTGTTGCC<br>GCATTAAAGACTAATCGTGTGACATCATCCGCGCTTCGCGCGGATGA<br>TTTTTCATGTCTGTGCCACGTATCTGTTGGCGGCATCCGCGCTTCGC<br>GCGGATGCCATGATTTTTCCCGACTGGAAAGAAAAACCACCCATCCG<br>CGCTTCGCGCGGATGGGATAACACGCAATTAATGTGAGTAATCAGCT<br>GACATCCGCGCTTCGCGCGGATGTCAAATGTTTAGTGCTCCTAAAGA<br>GACCGCCT | MHY-09 |
| 19 | TCAAATGTTTAGTGCTCCTAAAGAGACCGCCTTTCATCCGCGCTTCG<br>CGCGGATGAATTGTTTCCTCAATTCCTTTCAAAGCGTGGCCGCATCC<br>GCGCTTCGCGCGGATGCGTTCGGGTGACCAGATATTGATTTGCTTTA<br>GCACATCCGCGCTTCGCGCGGATGTGGCGCTTTGGCCGTCGTTTTA<br>CAAGCTATGAAGCATCCGCGCTTCGCGCGGATGCTTCCCAAAACCCT<br>GGCGTTACCCTTGTGAGCAGCATCCGCGCTTCGCGCGGATGCTAAC<br>CCTGGATTATATTGATGAACTAAATCCAACATCCGCGCTTCGCGCGG<br>ATGTTTACGTGATCAGGAATATGATGAAGTTGTCGGGCATCCGCGCT<br>TCGCGCGGATGCCTGAAAAGGTGGTTTCTTTGTTCATTAATAAAGCAT<br>CCGCGCTTCGCGCGGATGCTGGCTACGATACTGTCGTCGTCCATGG<br>CGAACACATCCGCGCTTCGCGCGGATGTGTTACTCGCACGGTTACG<br>ATGCGCGATCGCC | MHY-09 |
| 20 | TGTTACTCGCACGGTTACGATGCGCGATCGCCAGCATCCGCGCTTC<br>GCGCGGATGCTTGATGTGAAATGAATAATTCGCGGTAGGTTATCATC<br>CGCGCTTCGCGCGGATGATGAAATTAACTTGGTATTCAAAGATTTCT<br>GTATCATCCGCGCTTCGCGCGGATGATGTACTGTTATTGTTTCTCCC<br>GAACGTTAAACACATCCGCGCTTCGCGCGGATGTGTTTGCTCAAAAA<br>TTTAATGCGAGATGAAAGAGCATCCGCGCTTCGCGCGGATGCTAGTT<br>TTAACGTTTACAATTTAACGACGGGTTTCATCCGCGCTTCGCGCGGA<br>TGAAAAGGTATATTGACTCTTCTCAGTTTGTTTTTACATCCGCGCTTC<br>GCGCGGATGTAGTTATAATCGCTATGTTTTCAATAATTCAAAGCATCC<br>GCGCTTCGCGCGGATGCTTGCGATTTAATTAATAGCGACGATTGATT<br>TTTCATCCGCGCTTCGCGCGGATGAAATAAAGCATTAATTTATCAGCT<br>GATTCTCTCACATCCGCGCTTCGCGCGGATGTGAGGGTTATATTGAT<br>GGTGATTTTTGACATGTGCATCCGCGCTTCGCGCGGATGCACCAAAA | MHY-09 |

| | | |
|---|---|---|
| | CAACCCCTTCAGGTTGCACAGTAGCCATTTCTAACGCACCAGTCATACCAATAACTTAAGGGTAACTAGCCTCGCCGGCAATAGTTACCCTTATTATCAAGATAAGAAAGAAAAGGATTTTTCGCTACGCTCAAATCCTTTACAGAATTCGAGCTCCGTCGACAAG | |
| 21 | AAGCTTGCATGCCTGCAGGTCGACTCTAGAGAAGGGTAACTAGCCTCGCCGGCAATAGTTACCCTTATTATCAAGATAAGAAAGAAAAGGATTTTTGGATCCCCGGGTCCCATCCGCGCTTCGCGCGGATGGGAACAACTAATCAAAGAAGTATTCAAGTGATACCATCCGCGCTTCGCGCGGATGGTTATTACACTCAACCCTATCTCGTCCAAACTCGCATCCGCGCTTCGCGCGGATGCGCGCGTTCAGAAGGGTTCTATCTTCTTACGCAACATCCGCGCTTCGCGCGGATGTTTCAGGTGGCCGATTCATTAATGCCTCTCCCTCCATCCGCGCTTCGCGCGGATGGATCCTCTAGGTTCAGCAAGGTGAGAGGGTTTTCCATCCGCGCTTCGCGCGGATGGATATTTGAGAGTCGACCTGCAGGTACCCGGGCCCATCCGCGCTTCGCGCGGATGGGTGCCGGTACTCAAACTTTTAAACGCAAAATTCCATCCGCGCTTCGCGCGGATGGATAATGTAAAGCTGGCTGGAGTGCCAGAAGCAACATCCGCGCTTCGCGCGGATGTTGATGGCACTGTATATTCATCTGTGTAAAAGACCATCCGCGCTTCGCGCGGATGGTACTGTTGTTCCTATTGGTTAAAAATTATTT | MHY-10 |
| 22 | GTACTGTTGTTCCTATTGGTTAAAAATTATTTACATCCGCGCTTCGCGCGGATGTAGCCTTTTTATTCACTCACATATATTTACAGTTCATCCGCGCTTCGCGCGGATGAAGCAAGGGTAGATCTCTCAAAAATGACCTGACCCATCCGCGCTTCGCGCGGATGGGTCATAAAAAAAGTTTTCTCGCGAACTAAAATACATCCGCGCTTCGCGCGGATGTATATTTGTGTTTTTGGTACAACCTATTACAGAACATCCGCGCTTCGCGCGGATGTTCGCAGATACATGTTGGCGTTGTATGCCTCTGCCATCCGCGCTTCGCGCGGATGGCCTAAATATTGGGAATCAACTGTTCTACTCGCGCATCCGCGCTTCGCGCGGATGCGCGATATGAATGATAAGGAAAGATTAATACCCGCATCCGCGCTTCGCGCGGATGCGTTCTTGTTGAAGTCTTTCGGGCAATTAAAAGTCATCCGCGCTTCGCGCGGATGACTTCTTTAAAATTGTAGCTGGGTTGGTAAGAAACATCCGCGCTTCGCGCGGATGTTCAGGATTGCAAAAGCCTCTCGCCTGGCAAA | MHY-10 |
| 23 | TTCAGGATTGCAAAAGCCTCTCGCCTGGCAAACACATCCGCGCTTCGCGCGGATGTGACTGGTTCAATTACCCTCTGACTCTCTGATATCATCCGCGCTTCGCGCGGATGATTAGCGCATAATGAGCCAGTTCTCAACGTCCGACATCCGCGCTTCGCGCGGATGTCTGTACAGTTTGCTAACATACTGTGTATGTAAACATCCGCGCTTCGCGCGGATGTTTTCTACCCGTTCATCTGTCCTCGCGCCTGGTACATCCGCGCTTCGCGCGGATGTAGGTTGGTCTAATTCCCAAATGGTGGTGATTAACATCCGCGCTTCGCGCGGATGTTGCTGGCTGCCTTCGTAGTGGCATTTCGTTTGACATCCGCGCTTCGCGCGGATGTCGGCGCAGCGGTTCCGGTGGTGGGGCGGCTCCACATCCGCGCTTCGCGCGGATGTGAGGGAGACTATCGGTATCAAGCTGTCATTGAGCATCCGCGCTTCGCGCGGATGCTGGAAAGATCAAAAGCCATGTATAGTACACTGGCATCCGCGCTTCGCGCGGATGCCTGTATCACGACAAAACTTTAGAACTAACGTTGCATCCGCGCTTCGCGCGGATGCACCAAAACAACCCCTTCAGGTTGCACAGTAGCCATTTCTAACGCACCAGTCATACCAATAACTTAAGGGTAACTAGCCTCGCCGGCAATAGTTACCCTTATTATCAAGATAAGAAAGAAAAGGATTTTTCGCTACGCTCAAATCCTTTACAGAATTCGAGCTCCGTCGACAAG | MHY-10 |

**Table S4.** Copy number of msDNA in Ec67-WT and Ec67-17-nt insertion

| Ec67-WT | Weight (ng) | ssDNA copies | Copies of msDNA per Cell |
|---|---|---|---|
| MHY-01-1 | 0.83 | 2.43E+10 | 12.16 |
| MHY-01-2 | 1.95 | 5.73E+10 | 28.66 |
| MHY-01-3 | 0.48 | 1.40E+10 | 6.99 |
| | | **Ec67-WT average** | **15.93** |
| **Ec67-insertion** | **Weight (ng)** | **ssDNA copies** | **Copies of msDNA per Cell** |
| MHY-03-1 | 0.28 | 6.98E+09 | 3.49 |
| MHY-03-2 | 0.58 | 1.42E+10 | 7.12 |
| MHY-03-3 | 0.34 | 8.23E+10 | 4.12 |
| | | **Ec67-17-nt insertion Average** | **4.91** |

**Table S5.** Sequence information of 180 staples for the rectangular DNA origami tile.

| Name | Sequence (5'→ 3') |
|---|---|
| [02,04] | GAACGGTACAGAACAATATTACCGAATACCTA |
| [02,08] | TATAATCAGAACTCAAACTATCGGATGGATTA |
| [02,12] | TGTCCATCGATTAGTAATAACATCACACGACC |
| [02,20] | AGAGTCCATTTGATGGTGGTTCCGAGAGGCGG |
| [02,24] | GTCAAAGGACGCTGGTTTGCCCCATTTTTCTT |
| [04,04] | CATTTTGAATGCGCGAACTGATAGAACCACCA |
| [04,08] | TTTACATTAGACAATATTTTTGAACGGTCAGT |
| [04,12] | AGTAATAATTCTGACCTGAAAGCGACGCTGAG |
| [04,20] | TTTGCGTATTGCGTTGCGCTCACTGGGTACCG |
| [04,24] | TTCACCAGGGGTGCCTAATGAGTGTAGCTGTT |
| [06,04] | GCAGAAGATGAGGAAGGTTATCTATTAGAGCC |
| [06,08] | ATTAACACGGTCAGTTGGCAAATCTTTAGAAG |
| [06,12] | AGCCAGCAAACCTCAAATATCAAACAACTCGT |
| [06,20] | AGCTCGAAGGGTTTTCCCAGTCACTCTGGTGC |
| [06,24] | TCCTGTGTGTGCTGCAAGGCGATTGCCATTCA |
| [08,04] | GTCAATAGCTGATTATCAGATGATATTATACT |
| [08,08] | TATTAGACCCACCAGAAGGAGCGGCTACCATA |
| [08,12] | ATTAAATCGAGTAACATTATCATTGAAATAAA |
| [08,20] | CGGAAACCCGTGCATCTGCCAGTTTAATTCGC |
| [08,24] | GGCTGCGCGGTCACGTTGGTGTAGTCATCAAC |
| [10,04] | TCTGAATAAAGTTACAAAATCGCGCAAAAGAA |
| [10,08] | TCAAAATTAATAACGGATTCGCCTACAAAATT |
| [10,12] | GAAATTGCACAGTAACAGTACCTTAATTACCT |
| [10,20] | GTCTGGCCACGTTAATATTTTGTTGGTCATTG |
| [10,24] | ATTAAATGGAAGATTGTATAAGCAGAATCGAT |
| [12,04] | GATGATGAGCGATAGCTTAGATTAAAAATCAT |
| [12,08] | AATTACATATTAATTTTCCCTTAGCCTCCGGC |
| [12,12] | TTTTTAATAATAACCTTGCTTCTGGTAAATGC |
| [12,20] | CCTGAGAGAATATGATATTCAACCTAATACTT |
| [12,24] | GAACGGTATGAGAAAGGCCGGAGACGCAAGGA |
| [14,04] | AGGTCTGACGTGTGATAAATAAGGTACTAGAA |
| [14,08] | TTAGGTTGCTGACCTAAATTTAATTTATACAA |
| [14,12] | TGATGCAATTTTTCAAATATATTTCTCAACAG |
| [14,20] | TTGCGGGAGGCAAAGAATTAGCAAGTAGATTT |
| [14,24] | TAAAAATTTAGCATTAACATCCAACAAATGGT |
| [16,04] | AAAGCCTGAAAGTAATTCTGTCCAGCAGAACG |
| [16,08] | ATTCTTACTAATAAGAGAATATAAGTCCTGAA |
| [16,12] | TAGGGCTTATGTAATTTAGGCAGAATTTACGA |
| [16,20] | AGTTTGACCATGTTTTAAATATGCTAATTCGA |
| [16,24] | CAATAACCGCTTAATTGCTGAATAGCAAACTC |
| [18,04] | CGCCTGTTTTCATCGTAGGAATCAAGAAGGCT |
| [18,08] | CAAGAAAACACTCATCGAGAACAAGCGTTTTA |
| [18,12] | GCATGTAGCATTCCAAGAACGGGTTTTTGAAG |
| [18,20] | GCTTCAAACTGACTATTATAGTCAGCCAGAGG |
| [18,24] | CAACAGGTAATGACCATAAATCAAGGATAGCG |
| [20,04] | TATCCGGTAATAAACAGCCATATTTGTTTAAC |

| | |
|---|---|
| [20,08] | GCGAACCTCGTCTTTCCAGAGCCTTACAGAGA |
| [20,12] | CCTTAAATATTTTATCCTGAATCTCATTAGAC |
| [20,20] | GGGTAATAAGAGCAACACTATCATCGTTGGGA |
| [20,24] | TCCAATACCATAACGCCAAAAGGAAACTAACG |
| [22,04] | GTCAAAAAAAACAATGAAATAGCAAGTAAGCA |
| [22,08] | GAATAACAAGAATTGAGTTAAGCCGGAAACCG |
| [22,12] | GGGAGAATATTGAGCGCTAATATCCCCAAAAG |
| [22,20] | AGAAAAATTGAGATGGTTTAATTTGGCGCATA |
| [22,24] | GAACAACAGAGAAACACCAGAACGAATCTTGA |
| [24,04] | GATAGCCGTAAGTTTATTTTGTCAGCCAAAGA |
| [24,08] | AGGAAACGACATATAAAAGAAACGGAGGGAGG |
| [24,12] | AACTGGCACAAACGTAGAAAATACTTCATTAA |
| [24,20] | GGCTGGCTGAACGAGGCGCAGACGCGAAAGAG |
| [24,24] | CAAGAACCAAATCCGCGACCTGCTATCTTTGA |
| [26,04] | CAAAAGGGAACCATCGATAGCAGCCTTTAGCG |
| [26,08] | GAAGGTAAACCATTAGCAAGGCCGGCATTTTC |
| [26,12] | AGGTGAATTTAGAGCCAGCAAAATTGCCATCT |
| [26,20] | GCAAAAGAGGACTAAAGACTTTTTTGACAACA |
| [26,24] | CCCCCAGCAACGAGGGTAGCAACGTATTCGGT |
| [28,04] | TCAGACTGCCACCAGAACCACCACGGCAGGTC |
| [28,08] | GGTCATAGCGCCACCCTCAGAGCCAACAAATA |
| [28,12] | TTTCATAAACCGCCTCCCTCAGAGAAGCGCAG |
| [28,20] | ACCATCGCGCCTTTAATTGTATCGTTAGTAAA |
| [28,24] | CGCTGAGGTGAAAATCTCCAAAAATAAACAAC |
| [30,04] | AGACGATTCGTATAAACAGTTAATAAACATGA |
| [30,08] | AATCCTCATTAACGGGGTCAGTGCCAAGAGAA |
| [30,12] | TCTCTGAATTGATGATACAGGAGTTCAGTACC |
| [30,20] | TGAATTTTAAACTACAACGCCTGTCACCGTAC |
| [30,24] | TTTCAACATGTACCGTAACACTGATCAGAACC |
| [03,05] | GTAATATCCGCCAGAATCCTGAGAGTATAACG |
| [03,09] | GAGTAGAAGTGAGGCCACCGAGTAGAGCGGGC |
| [03,17] | AAAATCCCTGAGTGTTGTTCCAGTCGATTTAG |
| [03,21] | AAATCCTGCTATTAAAGAACGTGGAAGCACTA |
| [03,25] | AGCGGTCCGCGAAAAACCGTCTATCAAATCAA |
| [05,05] | AGTCTTTACGCTCAATCGTCTGAACCTTGCTG |
| [05,09] | CGTGGCACGGCAGATTCACCAGTCACTTGCCT |
| [05,17] | TCCAGTCGCGGCCAACGCGCGGGGAAATCGGC |
| [05,21] | CACATTAATTGGGCGCCAGGGTGGCAGGCGA |
| [05,25] | AAAGCCTGTGAGACGGGCAACAGCTTGCAGCA |
| [07,05] | AAAGGAATTAAAACAGAGGTGAGGTGGCTATT |
| [07,09] | CAATATCTCGCCTGCAACAGTGCCTAAGAATA |
| [07,17] | AAAACGACACTCTAGAGGATCCCCGCCCGCTT |
| [07,21] | TAACGCCATTCGTAATCATGGTCAAGCTAACT |
| [07,25] | AGGGGGATGAAATTGTTATCCGCTTAAAGTGT |
| [09,05] | TCATATTCATAATACATTTGAGGAAACAGTTG |
| [09,09] | CAAAGAAATTTACAAACAATTCGACCCTCAAT |
| [09,17] | CGACGACAGCTTTCCGGCACCGCTGACGTTGT |
| [09,21] | ATCGTAACAGGCAAAGCGCCATTCAAGTTGGG |
| [09,25] | ATGGGATAAACTGTTGGGAAGGGCCTGGCGAA |

| | |
|---|---|
| [11,05] | TGAATACCATGGAAGGGTTAGAACAATTATCA |
| [11,09] | GGAGAAACATTTGCACGTAAAACATTGCGGAA |
| [11,17] | CATTAAATGGAACGCCATCAAAAATGAGGGGA |
| [11,21] | AATTGTAATTCCTGTAGCCAGCTTATGGGCGC |
| [11,25] | AAAAACAGTGAGCGAGTAACAACCTGACCGTA |
| [13,05] | AAAACATAAACAAACATCAAGAAAGATTGCTT |
| [13,09] | CGCTATTATTAACAATTTCATTTGTTACATCG |
| [13,17] | TGATAAATTCTACAAAGGCTATCAAAAATTCG |
| [13,21] | TCACCATCTCTGGAGCAAACAAGAAATATTTA |
| [13,25] | CAAAAGGGATCGTAAAACTAGCATAAAGCCCC |
| [15,05] | ATACCGACGAGACTACCTTTTTAAAATCCTTG |
| [15,09] | TTCATCTTGGTTATATAACTATATTAAATCGT |
| [15,17] | AATAAAGCAAACATTATGACCCTGGTTCTAGC |
| [15,21] | ACAGGCAAGAAGCCTTTATTTCAACAGTCAAA |
| [15,25] | AATAGTAGTTTAGAACCCTCATATTAAAGATT |
| [17,05] | CAAAAGGTTTTAGTATCATATGCGGGTTTGAA |
| [17,09] | CGAGCCAGCAGTATAAAGCCAACGTAGTTAAT |
| [17,17] | TACGGTGTCCAATTCTGCGAACGAAATTAAGC |
| [17,21] | TAGCTCAACATTAGATACATTTCGTAAATCAT |
| [17,25] | GGCTTAGATGTTTAGCTATATTTTATTCTACT |
| [19,05] | TTTTTATTTATCAACAATAGATAAAGTACCGA |
| [19,09] | AAGTACCGATAATATCCCATCCTAGGCATTTT |
| [19,17] | GCGGATTGTTCAAATATCGCGTTTAACTAAAG |
| [19,21] | TCTTTACCGCGAACCAGACCGGAATAATGCTG |
| [19,25] | AAAACGAGCAGGATTAGAGAGTACTTGCGGAT |
| [21,05] | AGTTACAAATTCTAAGAACGCGAGGCAAGCCG |
| [21,09] | CTAACGAGCCCGACTTGCGGGAGGATTAAACC |
| [21,17] | TTTACCAGTTGCAAAAGAAGTTTTGAAGCAAA |
| [21,21] | GCATAGTAGTAAAATGTTTAGACTAAATCAGG |
| [21,25] | TGCAGATATGCGGAATCGTCATAACAGTTCAG |
| [23,05] | AGAGCAAGTGAAAATAGCAGCCTTAATTTGCC |
| [23,09] | AACCCACATAAAAACAGGGAAGCGTACCAACG |
| [23,17] | ATCATTGTATTATACCAGTCAGGAAACCCTCG |
| [23,21] | ATTGGGCTCTACGTTAATAAAACGATTACGAG |
| [23,25] | GCCCTGACTTATTACAGGTAGAAATCAACTAA |
| [25,05] | CCACGGAAAACAAAGTTACCAGAACAATAATA |
| [25,09] | AGGTGGCACAATAATAACGGAATAAGAGAGAT |
| [25,17] | TAAGGGAAAACGGTGTACAGACCACAACTTTA |
| [25,21] | CTTAGCCGGACCTTCATCAAGAGTAGTAGTAA |
| [25,25] | TTGTGTCGGGATATTCATTACCCATAAGGCTT |
| [27,05] | ACCAATGACGACATTCAACCGATTCAAAGACA |
| [27,09] | GCACCATTATATTGACGGAAATTAATACATAA |
| [27,17] | AGTTTCCAAGGCACCAACCTAAAAGTCAATCA |
| [27,21] | GGCTTTGAATACACTAAAACACTCCCATGTTA |
| [27,25] | AGCATCGGGATTATACCAAGCGCGCTGATAAA |
| [29,05] | CAGAGCCGTAGCGCGTTTTCATCGGAAACGTC |
| [29,09] | CTCAGAACCCCCCTTATTAGCGTTCACCAGTA |
| [29,17] | GCTTGCTTATAGTTGCGCCGACAACATGAGGA |
| [29,21] | CAAAAGGACCACGCATAACCGATAGCTACAGA |

| | |
|---|---|
| [29,25] | TTTCACGTCTTGCAGGGAGTTAAACGAAAGAC |
| [31,05] | ACAGTGCCGGCCTTGATATTCACAACCACCCT |
| [31,09] | AATAAGTTTTAAAGCCAGAATGGACCGCCACC |
| [31,17] | ACAGACAGGTCGTCTTTCCAGACGGTTTATCA |
| [31,21] | ACCAGTACCTGTATGGGATTTTGCAAAGGCTC |
| [31,25] | GGAACCCAGTTTCAGCGGAGTGAGAATAATTT |
| [02, 16] | GATAGGGTTTATAAATCAAAAGAAGTAGCAAT |
| [03, 13] | ACTTCTTTACGCAAATTAACCGTTTAGCCCGA |
| [18, 16] | CGAAAGACCATCAAAAAGATTAAGGGCTGTCT |
| [19, 13] | TTCCTTATAAACCAATCAATAATCAGGAAGCC |
| [04, 16] | TAATGAATGGAAACCTGTCGTGCCAACAGAGA |
| [05, 13] | TAGAACCCAAGGGACATTCTGGCCAGCTGCAT |
| [20, 16] | AGAGGCTTACGACGATAAAAACCATTTGCACC |
| [21, 13] | CAGCTACACAAGATTAGTTGCTATAAATAGCG |
| [06, 16] | GCAGGTCGGGCCAGTGCCAAGCTTAAGCATCA |
| [07, 13] | CCTTGCTGGCAAATGAAAAATCTAGCATGCCT |
| [22, 16] | ACTGGCTCGAATTACCTTATGCGAACAAAGTC |
| [23, 13] | AGAGGGTATAACTGAACACCCTGATTTTAAGA |
| [08, 16] | TCCAGCCAGTATCGGCCTCAGGAATTAATTTT |
| [09, 13] | AAAAGTTTCTTTGCCCGAACGTTAGATCGCAC |
| [24, 16] | GACAGATGCCGAACTGACCAACTTTTACGCAG |
| [25, 13] | TATGTTAGTGATTAAGACTCCTTATGAAAGAG |
| [10, 16] | AACCAATATTTTGTTAAATCAGCTAACGTCAG |
| [11, 13] | ATGAATATGTAGATTTTCAGGTTTCATTTTTT |
| [26, 16] | CACTACGATTAAACGGGTAAAATATTGAGCCA |
| [27, 13] | TTTGGGAATATCACCGTCACCGACCGTAATGC |
| [12, 16] | TTGAGAGATAATGCCGGAGAGGGTTCAATATA |
| [13, 13] | TGTGAGTGGGAAACAGTACATAAAAGCTATTT |
| [28, 16] | TGATACCGTCGAGGTGAATTTCTTCAGAGCCA |
| [29, 13] | CCACCGGATCAAAATCACCGGAACAAACAGCT |
| [14, 16] | TGTACCAACTCAGAGCATAAAGCTAAGAACGC |
| [15, 13] | GAGAAAACATCCAATCGCAAGACAAAATCGGT |
| [30, 16] | AAAGTTTTCCCTCATAGTTAGCGTGCGTCATA |
| [31, 13] | CATGGCTTTTTACCGTTCCAGTAAAACGATCT |
| [16, 16] | GTTGATTCCTGGAAGTTTCATTCCATTTAACA |
| [17, 13] | ACGCCAACAATTGAGAATCGCCATATATAACA |

**Table S6.** Sequence information of 14 linker strands used for cyclization of a tile into a tubule.

| Name | Sequence (5'→ 3') |
|---|---|
| [1,10] | GCTAGGGCATTAGCGGGGTTTTGCGTACTGGT |
| [1,14] | GAAAGGAAAAGTGCCGTCGAGAGGGTTGATAT |
| [1,18] | AGCTTGACCCCGGAATAGGTGTATAGCATTCC |
| [1,2] | AACAGGAGGGAACCTATTATTCTGGCCCCCTG |
| [1,22] | AATCGGAATTTAGTACCGCCACCCGTTTCGTC |
| [1,26] | GTTTTTTGCAGAACCGCCACCCTCGCCCAATA |
| [1,5] | TGCTTTCCAGAGGCTGAGACTCCTCTTGAGTA |
| [32,12] | AGGCGGATGGGAAGAAAGCGAAAGAAAGAGTC |
| [32,16] | AAGTATAGGGGGAAAGCCGGCGAACGTGGCGA |
| [32,20] | TCAGGAGGCCCTAAAGGGAGCCCCTTGGAACA |
| [32,24] | GCCACCCTGGGTCGAGGTGCCGTAACTCCAAC |
| [32,28] | CACCCTCATACGTGAACCATCACCCAGGGCGA |
| [32,4] | AAGTATTATCGTTAGAATCAGAGCTTAGACAG |
| [32,8] | GGATTAGGGCTGGCAAACGAGCACAGTGTTTT |

**References.**


1. Hsu, M.Y., Eagle, S.G., Inouye, M. & Inouye, S. Cell-free synthesis of the branched RNA-linked msDNA from retron-Ec67 of Escherichia coli. *Journal of Biological Chemistry* **267**, 13823-13829 (1992).
